# Task goals shape the relationship between decision and movement speed

**DOI:** 10.1101/2023.12.29.573524

**Authors:** Fanny Fievez, Ignasi Cos, Thomas Carsten, Gerard Derosiere, Alexandre Zénon, Julie Duque

**Author notes:** **CORRESPONDENCE TO: Fanny Fievez** Institute of Neuroscience Université catholique de Louvain 53, Avenue Mounier COSY- B1.53.04 B-1200 BRUSSELS Belgium Web: http://www.coactionslab.com.

## Abstract

The speed at which we move is linked to the speed at which we decide to make these movements. Yet, the principles guiding such relationship remain unclear: while some studies point towards a shared invigoration process boosting decision and movement speed jointly, others rather indicate a tradeoff between both levels of control, with slower movements accompanying faster decisions. Here, we aimed (1) at further investigating the existence of a shared invigoration process linking decision and movement and (2) at testing the hypothesis that such a link is masked when detrimental to the reward rate. To this aim, we tested 62 subjects who performed the tokens task in two experiments (separate sessions): Experiment 1 evaluated how changing decision speed affects movement speed while Experiment 2 assessed how changing movement speed affects decision speed. In the latter experiment, subjects were either encouraged to favor decision speed (fast decision group) or decision accuracy (slow decision group). Various mixed model analyses revealed a coregulation of decision (urgency) and movement speed in Experiment 1 and in the fast decision group of Experiment 2, but not in the slow decision group despite the fact that these same subjects displayed a coregulation effect in Experiment 1. Altogether, our findings support the idea that coregulation occurs as a default mode but that this form of control is diminished or supplanted by a tradeoff relationship, contingent on reward rate maximization. Drawing from these behavioral observations, we propose that multiple processes contribute to shaping the speed of decisions and movements.

**New & NoteWorhty:** The principles guiding the relationship between decision and movement speed are unclear. In the present behavioral study involving two experiments conducted on 62 human subjects, we report findings indicating a relationship that varies as a function of the task goals. Coregulation emerges as a default mode of control that fades when detrimental to the reward rate, possibly due to the influence of other processes that can selectively shape the speed of our decisions and movements.

## Introduction

Every day of our life, we make decisions to interact with our environment in a goal-directed manner. Several models have been developed to explain how we select actions by describing decision making as a process of noisy accumulation of evidence: during deliberation, sensory evidence is accumulated until the total evidence in favor of one of the potential actions reaches a certain threshold, at which point the decision is made and an action is initiated (Gold & Shadlen, 2007; Hanks et al., 2014; Ratcliff et al., 2016). In the context of motor behavior, such accumulation directly converts into an increase of neural activity in the motor cortex, from a starting point to a consistent decision threshold (Cisek & Thura, 2022; Kelly et al., 2021; Ratcliff & Smith, 2004). In these models, the time taken to select a motor act is a direct function of the speed with which motor activity grows (based on evidence) and of the amount of activity required to reach the threshold, which sets the accuracy criterion.

Decisions can be made even when evidence is low or absent, suggesting that the accuracy criterion can be reduced if necessary. For instance, if we unexpectedly reach a junction when driving a car, we must rapidly decide whether to turn left or right, regardless of readiness and evidence, increasing the risk of an inappropriate choice. Past studies have suggested that this decrease in accuracy criterion can be implemented by an evidence-independent rising “urgency signal” pushing motor activity close to decision threshold, which is mathematically equivalent to a lowering of the latter (Churchland et al., 2008; Cisek et al., 2009; Ditterich, 2006; Murphy et al., 2016; O’Connell et al., 2018; Thura, 2020). Computationally, this urgency signal is typically modelled as increasing linearly over time (Cisek et al., 2009; Derosiere et al., 2019, 2021, 2022; Kelly et al., 2021) it includes an intercept, which reflects the degree to which motor activity is upregulated already at the onset of the decision process (i.e., context-sensitive effect), and a slope, which reflects how fast motor activity will be pushed towards the decision threshold as time elapses (i.e., time-sensitive effect).

Reward plays an important role in animal behavior and “reward rate” maximization is a prominent principle in the field of decision making (Lemon, 1991; Shadmehr et al., 2019). More precisely, the reward rate is the sum of all rewards acquired through our actions, minus the costs implied over the total time spent (Carland et al., 2019; Yoon et al., 2018). Critically, the urgency signal described in models of decision making is typically considered as useful for maximizing the reward rate (Carland et al., 2019; Charnov, 1976; Thura et al., 2012). That is, by its context-sensitive effect, the urgency signal can shorten the time invested to obtain the reward. In addition, by its time-sensitive effect, it allows to make decisions without waiting indefinitely, even when there is only little or no evidence.

Several observations in both human and non-human primates indicate that the time one takes to decide impacts the speed of the movement implementing the decision (Churchland et al., 2008; Thura et al., 2014; Drugowitsch et al., 2012). That is, faster decisions are typically followed by quicker movements. Interestingly, the reverse relationship has been observed too: one study in our lab has revealed that when the context requires movements to be performed in a shorter time period, the decisions leading to them arise faster, compared to contexts in which subjects have more time to perform the same movement (Carsten et al., 2023). Altogether, these findings suggest that the urgency signal implemented in decision making models may not solely operate at this restricted level but may shape behavior in a global manner, coregulating the speed of both decisions and movements (Carsten et al., 2023; Thura, 2020). This “coregulation hypothesis” is consistent with the view that the urgency signal serves to maximize reward rate, as in most situations a speeding up of both decision and movement will shorten the time required to obtain a reward (Shadmehr et al., 2019). Also consistent with this hypothesis is the fact that some studies have reported a time-sensitive effect of decision urgency on movement speed: that is, within a given context, movements that are initiated following a longer deliberation are typically faster than movements associated with earlier decisions, reflecting well the linear increase in urgency over time (Thura et al., 2014; Thura, 2020).

The “coregulation hypothesis” has received systematic support from studies investigating the impact of decision urgency on movement speed (Carsten et al., 2023;Thura et al., 2014; Thura, 2020). By contrast, studies that have addressed this relationship inversely, by looking at the impact of movement speed on the pattern of decision speed, have reported mixed findings (Carsten et al., 2023; Kita et al.,2023; Reynaud et al., 2020; Saleri Lunazzi et al., 2021). As mentioned above, we recently observed changes in decision speed in relation to context-dependent adjustments in movement speed that are consistent with the coregulation hypothesis (Carsten et al., 2023). It is interesting to note that in the latter study, the increase in decision speed when subjects had to perform faster movements occurred even when there was no possibility of saving time by doing so; that is, making faster decisions did not allow to shorten the trial/block duration. This led us to think that coregulation of decision and movement may be a ubiquitous feature of human behavior, one that occurs by default even when not required, arising from a neural organization that has developed because most real-life (urgent) situations require rapid actions (i.e., including fast decisions and fast movements). Yet, in their recent work, Thura and his team have reported a slowdown of decision speed (rather than a speedup) in contexts requiring subjects to perform faster movements (Reynaud et al., 2020; Saleri Lunazzi et al., 2021), reminiscent of a tradeoff rather than a coregulation effect. Yet importantly, this bunch of work used an experimental design in which subjects knew that a block would only end after a fixed number of correct decisions. Hence, here speeding up decisions was detrimental to the reward rate, as the gain of time on a trial basis would have led to lengthening the experiment, given the decline in accuracy that typically accompanies faster decisions.

Based on these collective findings, it appears that if an urgency mechanism exists in the brain, it must interact with other processes that provide human beings with a flexible control of decision and movement for adaptive behavior according to the task goals, as previously suggested (Saleri Lunazzi et al., 2021) but never tested directly. Here, we addressed this idea in two behavioral experiments on healthy young human participants who performed a variant of the Tokens task in different contexts. Specifically, we tested the prediction that decision and movement durations would be linked by default in this task but that this coregulation would disappear in the same subjects when detrimental in terms of reward rate, as a function of the context in which they perform the task.

## Methods

### Participants and ethical statement

A total of 62 healthy human volunteers were recruited for this study, but due to technical issues the data were processed on 56 subjects (30 women, 24.6 ± 3.4 years old). All participants were right-handed according to the Edinburgh Questionnaire (Oldfield, 1971) and had normal or corrected-to-normal vision. None of them had any neurological disorder or history of psychiatric illness or drug or alcohol abuse; and no one was following any clinical treatment that could have influenced performance. The protocol was approved by the Ethics Committee of the Université catholique de Louvain (UCLouvain), Brussels, Belgium (approval number: 2018/22MAI/219) and adhered to the principles expressed in the Declaration of Helsinki. Participants were financially compensated for their participation and provided written informed consent.

### Setup and task

Experiments were conducted in a quiet and dimly lit room. Participants were seated in front of a 21-inches cathode ray tube computer screen, placed at 60 cm from the participant’s eyes and used to display stimuli during the task. The display was gamma-corrected, and its refresh rate was set at 75 Hz. Participants’ forearms were positioned in a neutral position (i.e., at 0 degree of pronation and supination) on a Canadian board, used to standardize and fix the resting position of the index fingers with elastic bands. Moreover, two lasers (targeting the tip of each index finger) were used to indicate the resting position (i.e., 0 degree of index flexion) and the minimum amplitude of the left or right index finger movement required in the task (i.e., 30 degrees of flexion, see Fig. 1.A and below for more details).

**Figure 1.**
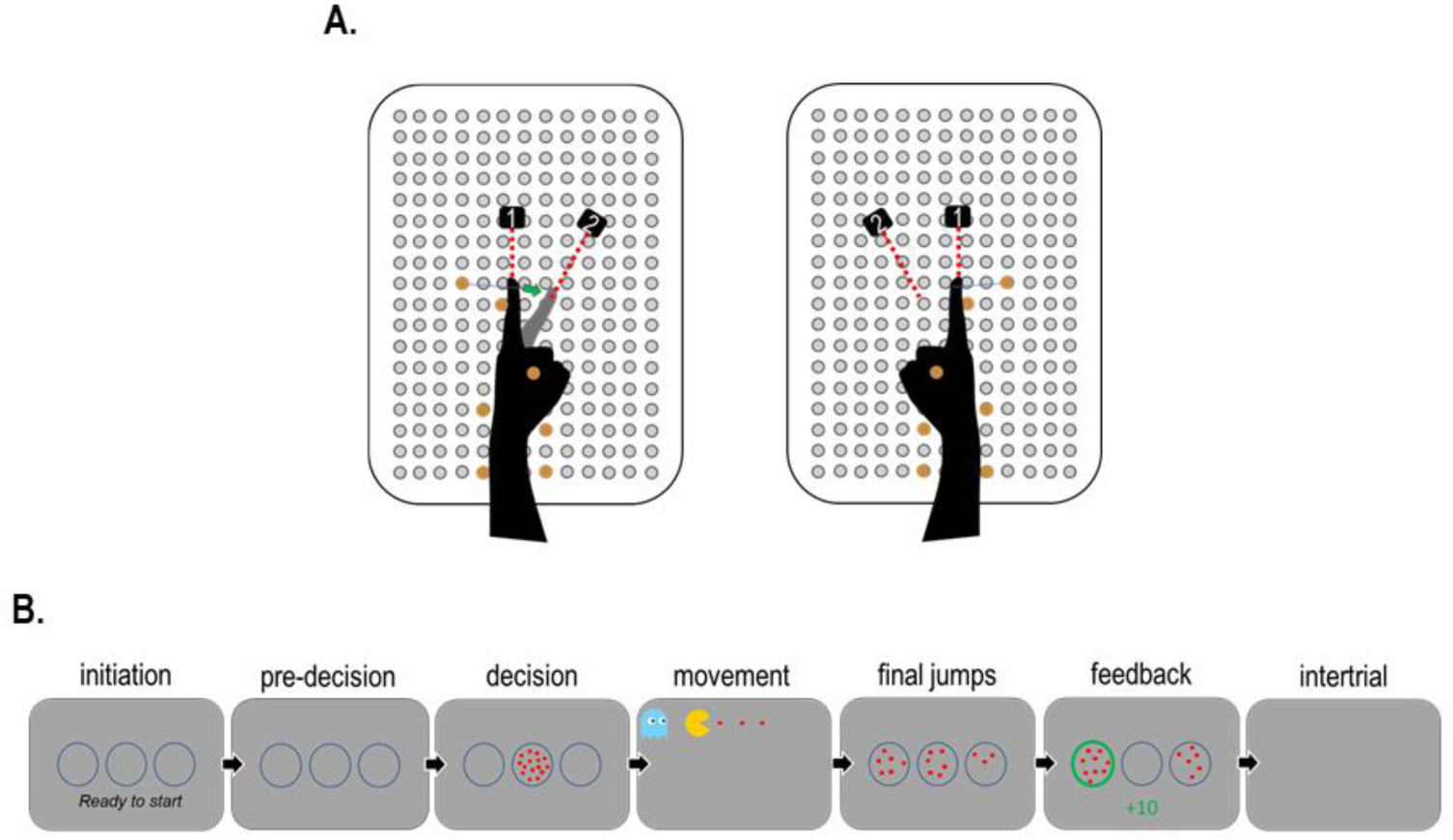
Experimental material. **A. Setup.** Participant forearms were positioned in a neutral position on a Canadian board, with elastic bands maintaining the index fingers in a resting position. Two lasers (targeting the tip of each index finger) were used to indicate the resting position (i.e., 0 degree of flexion; laser 1) and the minimum amplitude (i.e., 30 degrees of flexion; laser 2) of the left or right index finger movement required in the task. **B. Task.** Each trial starts with an initiation phase during which participants have to perform a bilateral flexion of index fingers to signal their readiness. When index fingers return to their resting position, the 3 circles remain empty for a variable pre-decision period (150-300 ms). Then, 15 tokens appear in the central circle and start jumping one by one every 200 ms. Subjects are required to determine, as soon as they feel ready, which lateral circle will end up with the largest number of tokens (left one in this example), and to report their decision with a left or right index finger movement triggering a pacman animation (more details can be found in the main text). Once the movement phase is over, the central circle finishes to empty with the final token jumps. The trial ends with a feedback (fully successful trial in this example; more in the main text) followed by a blank screen of variable intertrial duration (1800-2000 ms).

We used a variant of the Tokens task (Cisek et al., 2009;Thura et al., 2014) which has already been exploited in various ways in several past studies of our lab (Derosiere et al., 2019, 2022; Fievez et al., 2022) The current version of the task was implemented with Matlab 2016 (The Mathworks, Natick, Massachusetts, USA) and the Cogent 2000 toolbox (Functional Imaging Laboratory, Laboratory of Neurobiology and Institute of Cognitive Neuroscience at the Welcome Department of Imaging Neuroscience, London, UK).

#### Sequence of events in a trial

The overall sequence of events for each trial of our task is depicted in Figure 1.B. Each trial starts with the appearance of 3 empty blue circles (4 cm diameter each), placed on a horizontal axis, with the sentence “Ready to start” displayed below them. During this initiation phase, subjects are required to perform a bilateral flexion of index fingers to indicate that they are ready to start the trial. When index fingers return to their resting position, the 3 circles remain empty for a short pre-decision phase of random duration (150-300 ms). The decision phase then starts with the appearance of 15 randomly arranged tokens (0.3 cm diameter) in the central circle that start to jump one by one from the center to one of the 2 lateral circles, every 200 ms. The task of the subjects is to determine which of the two lateral circles will end up with the largest number of tokens and to report their decision with a left or right index finger movement. The movement phase involves a pacman animation that will be described below in more details. Once this phase is over, the final tokens continue to jump until the central circle is empty (i.e., until the 15^th^ token jump; Jump_-15_). Note that subjects are told that they are expected to respond before Jump_-15_, as soon as they feel sufficiently confident. The trial always ends with a feedback (see below for more details) which is followed by a blank screen during a variable intertrial interval (1800-2000 ms).

#### Trial types and success probability

The difficulty of the decision in each trial of the task depends on the dispersion of tokens along the jumps, which could point more or less obviously to one of the lateral circles. This pattern of token distribution and the degree to which it affects success of the subjects can be formalized with what is called the “success probability”. That is, for each trial *i*, and at each moment in time, we can define the success probability p_i_(t) associated with choosing a response (left or right). If at a moment, the left (L) circle contains *NL* tokens, the right (R) once contains *NR* tokens, and *NC* tokens remain in the central (C) circle, then the probability that the left response is ultimately the correct one (i.e., the success probability of guessing left) is described as follows:

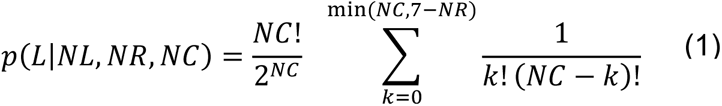

Calculating this quantity for the 15 token jumps allowed us to characterize the temporal profile of success probability *p_i_(t)* for each trial. As such, as far as the participants knew, the individual token jumps and the correct choice were completely random. However, we interspersed distinct trial types within the full sequence of trials. First, in 60% of trials, the *p_i_(t)* remained between 0.33 and 0.66 up to Jump_-8_, that is, the initial token jumps were balanced between the lateral circles, keeping the *p_i_(t)* close to 0.5 until late in these “ambiguous” trials. As such, in ambiguous trials, the tokens jumped alternatively to the correct and incorrect lateral circles until Jump_-8_, such that the number of tokens was equal in both lateral circles after each even jump (i.e., after 2, 4, 6 and 8 jumps) and a difference of one token was present after each odd jump (i.e., after 1,3,5 and 7 jumps). Second, in 20% of trials, the *p_i_(t)* was above 0.7 after Jump_-3_ and above 0.8 after Jump_-5_, that is, the initial jumps consistently favored the correct choice in these “obvious” trials. In the remaining 20% of trials, the *p_i_(t)* was below 0.4 after Jump_-3_, that is, the initial jumps favored the incorrect choice and the following ones favored the correct choice in these “misleading” trials.

#### Sensory evidence and SumLogLR

As in previous studies (Cisek et al., 2009; David Thura et al., 2014; Derosiere et al., 2022), we assumed that participants would estimate the level of evidence based on the number of tokens that have already jumped into the two side circles rather than by calculating the probability of success. This estimation can be computed as a first-order approximation of the real probability function after each jump (see Eq. 1), called the sum of log-likelihood ratios (SumLogLR):

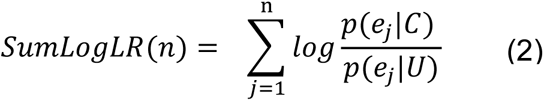

In this equation*, p(ej |C)* is the likelihood of a token event *ej* (a token favoring either the chosen or the unchosen response) during trials in which the chosen response C is correct, and *p(ej |U)* is the likelihood of e_j_ during trials in which the unchosen response U is correct. The SumLogLR is proportional to the difference between the number of tokens that favored each of the 2 possible choices (i.e., that moved toward each lateral circle) at any given time.

#### Index finger movement and pacman animation

As explained in more detail below in the “Experimental design” section, testing our predictions required us to characterize and/or to manipulate the movement features of index finger responses. To allow us to do so, we asked the subjects to provide their response, not with a single finger movement as we usually do, but with a tapping movement involving a repetition of 4 index finger flexions, with the left or right hand depending on their decision in the Tokens task. Moreover, this tapping movement was associated with a pacman animation. That is, as soon as the first tap was completed (i.e., flexion of at least 30 degrees, as indicated by the laser device), a pacman appeared together with 3 additional tokens in front of it, on a horizontal axis, as shown on Figure 1.B. Subjects knew that each further tap would allow the pacman to “eat” one of these supplementary tokens. The pacman was presented together with a ghost which will be described in the “Experimental Design” section, as its features depended on the experimental condition the subjects were in. Notably, the pacman animation often ended with a blank screen to cover the minimum 3000 ms period of the movement phase: as such, completing the tapping movement faster never allowed to shorten the trial duration, as in Carsten et al, 2023. Finally, note also that trials in which subjects did not provide a response before Jump_-15_ did not involve any pacman animation; in this case participants had instead to remain still in front of a blank screen for a period corresponding to the movement phase duration.

#### Trial feedback and score calculation

As a feedback at the end of the trial, the chosen circle turned either green or red, depending on whether the choice was correct or incorrect, respectively. The feedback also included a numerical score presented below the central circle, which depended on both the decision and movement requirements of the task. Subjects got +5 points for a correct decision and another +5 points for a correct tapping movement (see below for specific movement requirements). Hence, fully successful trials were worth +10 points (i.e., +5+5), as illustrated on Figure 1.B. Then, any failure, whether at the level of the decision or the movement, was penalized by -2 points. Hence, a partially successful trial was worth +3 points (i.e., +5-2 or -2+5), while a fully failed trial led to - 4 points (i.e., -2-2). Finally, in the absence of response before Jump_-15_ (i.e., a no-response trial), the displayed score corresponded to 0 point and came along with a ‘Time Out” message at the bottom of the screen. All trial scores were summed up and displayed at the end of each block, providing a global feedback on the block to maintain motivation of participants.

### Experimental Design

Participants performed two experiments as part of the current study. These experiments occurred on separate days (average interval of 9 ± 8 days) and in a counterbalanced order. In Experiment 1, we designed the task such that, in separate blocks, subjects were either encouraged to make slow or fast decisions. The purpose here was to further investigate the existence of a shared invigoration process linking decision and movement by looking at how contextual changes in decision speed would influence movement speed. Inversely, in Experiment 2, we aimed at testing the reverse relationship, thus how contextual changes to movement speed (slow or fast) influence decision speed. In addition, to test the hypothesis that such a link is masked when detrimental to the reward rate, we separated subjects in two groups who performed the slow and fast movement blocks in a context that either favored fast (fast decision group) or slow decisions (slow decision group). As such, we predicted that fast movement blocks would involve faster decisions (as compared to slow movement blocks) in the fast decision group (where coregulation is beneficial) but not in the slow decision group (where coregulation is detrimental). The specific features of these two experiments are described in the sections below.

#### Experiment 1

All subjects (n=56) took part in Experiment 1. This experiment required participants to perform the Tokens task in two different block types, where a small adjustment to one trial event (the final jumps phase) allowed us to promote either slow (accurate) or fast decisions (see Fig. 2.A). That is, in the “slow decision” blocks, tokens kept on jumping every 200 ms during the final jump phase, as in the decision phase, until the central circle was empty. In contrast, in the “fast decision” blocks, final tokens jumped much faster (i.e., every 50 ms), such that deciding faster allowed subjects to finish the trial earlier. Because blocks had a fixed duration of 265 sec, deciding earlier in fast decision blocks allowed subjects to perform more trials and thus to accumulate more points. This was not the case in the slow decision blocks, where it was a better strategy to decide later and reach a higher accuracy, given the fixed number of trials (26) that they would be able to do over the same block duration (always about 265 sec). All of this was made explicit to the subjects to enhance the change in decision policy between the two block types. Finally, as mentioned above, the pacman animation during the movement phase also involved a ghost which, in this experiment, always chased the pacman at a fixed speed established during pilot tests. Subjects knew that they had to avoid the ghost and would get a negative feedback (-2 points, as explained in the feedback section above) if they tapped so slowly that they got caught, in which case the pacman turned into a skull. Yet this happened very rarely because the imposed speed was easy to attain, as it was just there to exhort subjects to do their tapping movement in a row, as opposed to encourage them to go as fast as possible. That is, subjects were able to escape the ghost on most trials, regardless of the block type (see Fig. S1 in supplementary materials).

**Figure 2.**
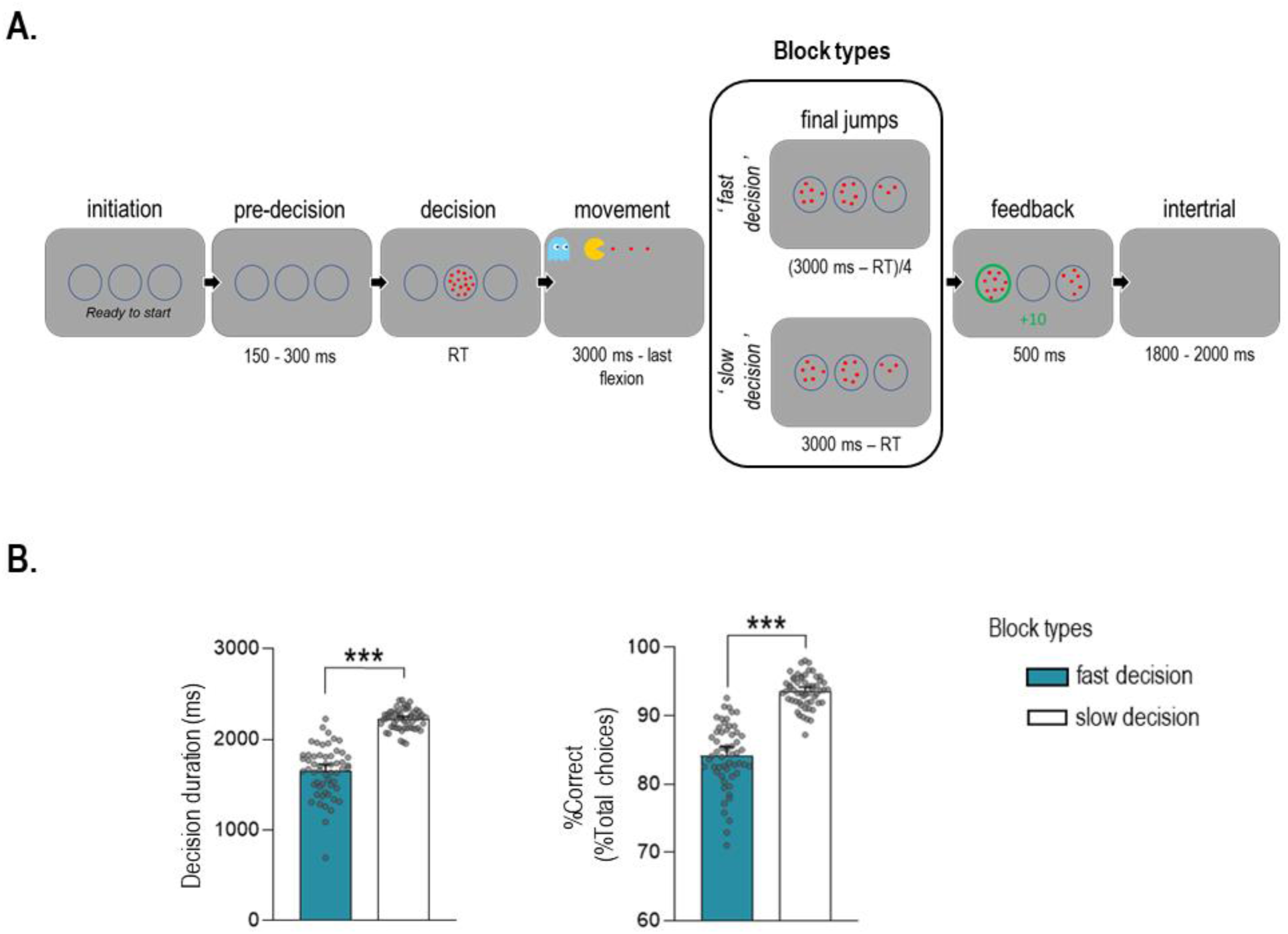
Protocol of Experiment 1. **A. Task.** In this experiment, participants performed two block types that differed according to the speed at which the final tokens jumped in the lateral circles at the end of the trial (after the finger tapping movement). That is, fast final jumps (every 50 ms) led participants to prioritize decision speed (fast decision blocks) while slow jumps (every 200 ms) led subjects to rather prioritize decision accuracy (slow decision blocks). **B. Manipulation check.** Participants adjusted their decision policy depending on the type of block by making faster (shorter decision duration) and less accurate (lower %Correct) decisions in fast (blue bars) relative to slow (white bars) decision blocks. Error bars represent SE. ***p < 0.001.

#### Experiment 2

All subjects also performed Experiment 2 but due to a lack of data after cleaning, 2 subjects had to be excluded from the analysis (n=54; 32 Women, 24.5 ± 3.3 years old). As shown on Figure 3.A, in this experiment again there were two different block types, which here differed according to the movement speed requirement. That is, subjects were explicitly asked to report their decision in the Tokens task with a tapping movement that was either fast, in the “fast movement” blocks, or slow, in the “slow movement” blocks. The exact speed that subjects had to adopt in both block types depended on their spontaneous speed. In the fast movement blocks, subjects had to tap at a speed that was twice the spontaneous speed (± 20%), while the speed in the slow movement blocks had to be equal to the spontaneous speed or to any value below it. The spontaneous tapping speed was calculated during a calibration phase at the beginning of the experiment, by asking subjects to perform a block of 15 trials of the Tokens task in a neutral condition, with no specific instruction regarding decision or movement speed. Once the average spontaneous speed was determined based on these trials, the participants were then specifically trained to perform fast tapping movements at the required speed (i.e., twice the spontaneous speed) during 4 blocks for each index finger. In these blocks, subjects repetitively tapped their index finger and each of these movements was followed by a feedback indicating if the tapping was “ok”, “too fast” (>20% above the targeted speed) or “too slow” (<20% below the targeted speed). A block ended as soon as subjects had completed 9 tapping movements at the targeted speed. There was no training for the speed of slow movement blocks as this speed was the one obtained spontaneously.

**Figure 3.**
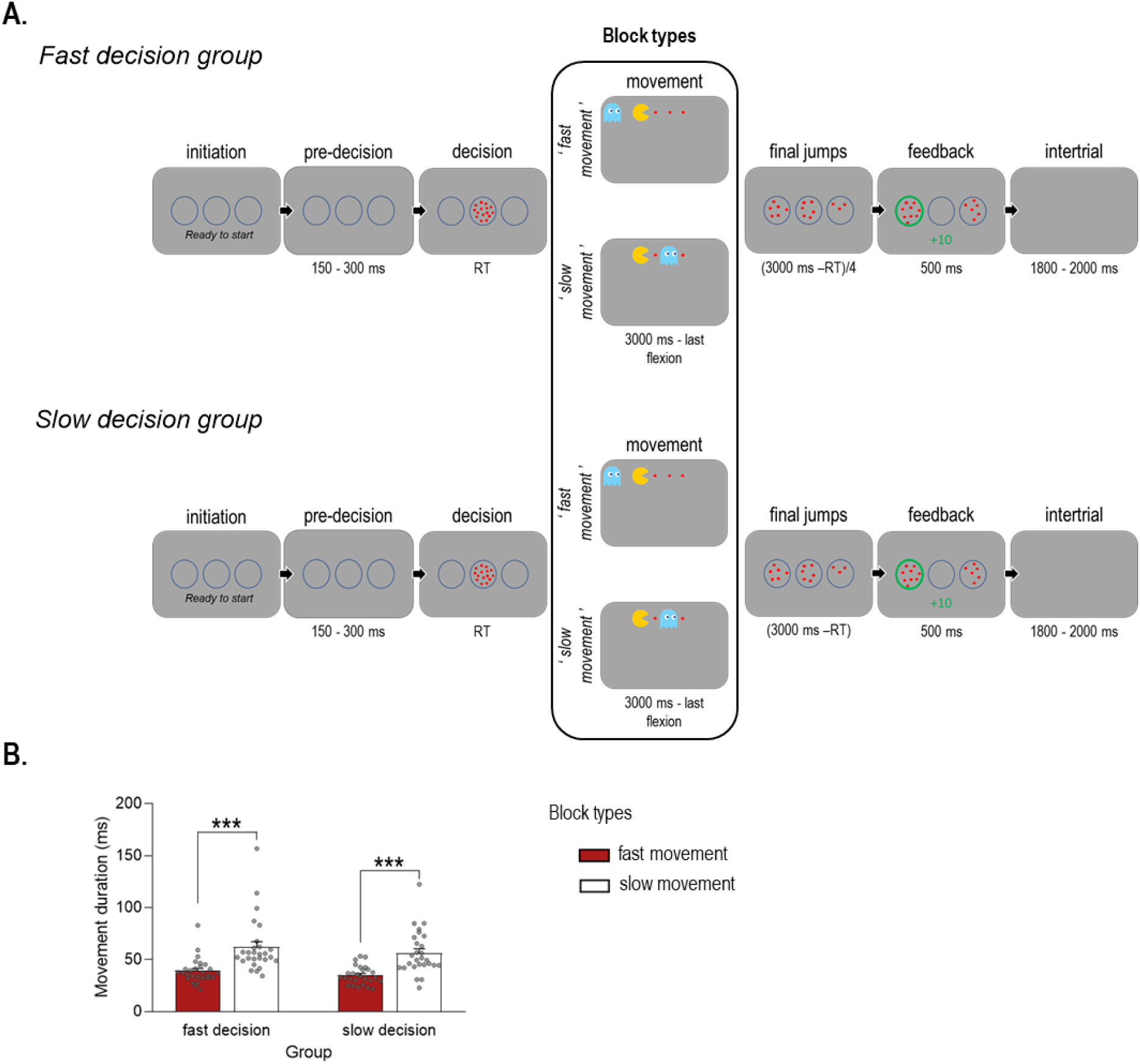
Protocol of Experiment 2. **A. Task.** In this experiment, participants performed two block types that differed according to the speed at which they had to perform the finger tapping movement; that is, at twice the spontaneous speed (fast movement blocks: pacman chased by ghost) or at about the spontaneous speed (slow movement blocks; pacman preceded by ghost). Moreover, subjects were split into two experimental groups according to whether they performed the version of the task that led them to prioritize decision speed due to fast (every 50 ms) final jumps (fast decision group, upper panel), or to prioritize decision accuracy due to slow (every 200 ms) final jumps (slow decision group, lower panel). **B. Manipulation check.** As required by the instruction in this Experiment, participants displayed shorter (faster) tapping movements in the fast (red bar) than slow (white bar) movement blocks, regardless of whether they belong to the fast or slow decision group. Error bars represent SE. ***p < 0.001.

As in Experiment 1, tapping movements in the Tokens task were performed in relation to a pacman (and ghost) animation, where each tap provided the opportunity to “eat” an additional token placed on a horizontal axis. Yet, here the animation served the further purpose to control for the movement speed instruction in each block type. First, to provide a guide to subjects, the ghost here was either moving behind the pacman, chasing it, in the fast movement blocks, while it was preceding the pacman, in the slow movement blocks (see Fig. 3.A). Given the continuous requirement to avoid a collision with the ghost, this feature reminded the subjects to tap rapidly (to escape the ghost) in the fast movement blocks and incited them to tap tranquilly (to avoid bumping into the ghost) in the slow movement blocks.

Importantly, a specificity of Experiment 2 is that we separated subjects in two groups who performed the slow and fast movement blocks in slightly different contexts, due to a small difference in the Tokens task. That is, the final tokens either jumped every 50 ms or every 200 ms, allowing subjects to shorten the trial duration by deciding faster in the group performing the task in the former condition (fast decision group) but not in the group using the latter version (slow decision group). Here, the number of trials was always fixed such that shortening the trial duration never allowed to do more trials. Yet, by responding faster, subjects could save time and finish the experiment earlier. Even if these specificities remained implicit in this experiment, we expected that they would lead subjects to adjust their decision speed such that it would be generally faster in the fast decision group (n=27; 15 women, 25.1 ± 3.6 years old), and generally slower in the slow decision group (n=27; 17 women, 23.9 ± 2.9 years old). With these two groups, we aimed at testing the hypothesis that a default mode of coregulation can be observed when it naturally serves the reward rate; but it would disappear in situations where it is detrimental to the reward rate. As such, we predicted that tapping faster in the fast movement blocks will translate into faster decisions, compared to the decisions in the slow movement blocks, but that this effect will only be observed in the group of subjects incited to make fast decisions but not in the slow decision group where subjects should better favor decision accuracy. Importantly, each subject completed the short version of the UPPS-P Impulsive Behavior (UPPS) scale (Eben et al., 2020). We also evaluated their ability to follow movement instructions in order to make sure that the repartition of participants in the two groups was not creating a bias in our results. The analyses showed no significant difference between the two groups of participants, considering their UPPS scores (see Table S1 in supplementary materials) or their movement performance (see Fig. S2 in supplementary materials).

#### Time course of the experiments

Experiments 1 and 2 had a largely similar time course. They both started with a familiarization phase during which subjects performed a neutral block of 10 trials to become acquainted with the basic features of the Tokens task. Subjects were then informed about the two types of blocks they would realize in the experiment. This led, after a phase of calibration (in Experiment 2 only to establish the fast tapping speed), to a training phase during which subjects practiced for 4 blocks (2 blocks of each type), each involving 15 trials (or a few more in fast decision blocks of Experiment 1). The actual experiment then started and involved a total of 18 blocks (9 blocks of each type), each involving 26 trials (or a few more in fast decision blocks of Experiment 1). Each block lasted on average 4,36 minutes and a break of 2 to 5 minutes was provided between blocks.

Both experiments then ended with blocks consisting of a choice reaction time (CRT) version of the Tokens task. The task in these blocks (always 40 trials) was the same as in the main experiment: subjects had to respond with a left or right index finger tapping movement according to token jumps but here the 15 tokens jumped all at once in the left or right circle. Experiment 1 entailed 1 CRT block with no restriction on movement speed (i.e., “free” movements), whereas Experiment 2 entailed 1 CRT block for each tapping speed (slow or fast) condition, considering that tapping speed may impact reaction time (RT).

### Computational approach

In our work we simulated the decision data by using an implementation of the urgency gating model (UGM; Cisek et al., 2009; Thura et al., 2014), in which evidence is multiplied by a linearly increasing urgency, as follows:

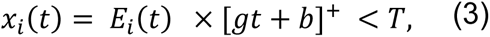

where *x*_*i*_(*t*) represents the activity of motor neurons at time *t* in trial *i*, obtained by multiplying the momentary evidence *E*_*i*_(*t*) by an urgency signal. *g* and *b* are the slope and the y-intercept of the urgency signal, and [ ]^+^ denotes half-wave rectification (which set all negative values to zero). To estimate the level of evidence accumulation, we used the computation of the SumLogLR (see above, equation 2) that we low-pass filtered for dealing with intra-trial stimulus noise when calculating evidence such as the evidence accumulation. The accumulation of evidence can thus be expressed as follows:

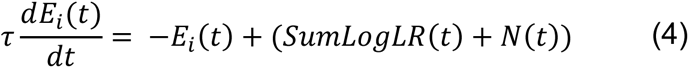

The Term *N*(*t*) represents Gaussian noise with a mean of zero and a SD of 0.7 and, we used a linear differential equation with a time constant τ = 200 ms as low-pass filter (Cisek et al., 2009).

The model has three parameters: *g*, *b* and *T*. To fit the data, we performed an exhaustive parameter grid search, with *g* ranging from 1 x 10^-7^ to 4, *b* from - 1 to 3 and *T* from 1 to 3. This was performed separately for each subject and each block condition, and the quality of fit was assessed by using the mean square-error between data distributions of the model and the real data distributions for all decision durations in the interval between 600 and 3000 ms. Importantly, the threshold was not fixed between participants (i.e., ranged from 1 to 3) but was fixed across block conditions (i.e., for fast and slow conditions of the same experiment) to be able to compare the urgency parameters between them. After determining the best fitting parameters for each dataset, we generated a subsequent grid search from the average of the gain and slope of all subjects to determine the best pairs of parameters for a given threshold. This procedure was repeated up to 100 times to ensure parameter optimality convergence, we computed the mean shape (linear function based on g and b parameters) of the resulting urgency functions.

### Data analyses

Behavioral data were collected by means of custom Matlab scripts (MathWorks, Natick, Massachusetts, USA). Statistical analyses were performed using JASP (Wagenmakers et al., 2018) for repeated-measures analyses of variance (ANOVA_RM_), t-tests and Spearman’s correlations, and using the package gamlj in jamovi (ŞAHİN & AYBEK, 2020) for linear mixed models.

#### Data processing and definition of endpoint measures

In general, trials with poor performance in the decision phase and/or in the movement phase of the Tokens task were excluded in both experiments. This concerned trials with no response before Jump_-15_ or with an anticipated response (before Jump_-3_). This also involved removing trials where the pacman was caught by the ghost and, for Experiment 2, also trials in which the participants did not move at the required speed. Finally, the laser sometimes failed to report the index finger movements and these trials were removed too. As a result, overall, 24% of trials were removed for the analyses (17% for Experiments 1 and 31% for Experiment 2). The same procedure was followed to clean the data in the CRT task, which led to an overall removal of 11% of trials (1% and 20% for Experiments 1 and 2, respectively).

The endpoint measures were the same for Experiment 1 and 2. To characterize the speed at which subjects made their choice, we considered the time they took to decide. This *decision duration* was computed as the average RT in the Tokens task (computed separately for fast and slow blocks with all trial types pooled together) minus the average RT in the corresponding CRT block (Ferrucci et al., 2021). It should be noted that the majority of studies have used the RT in simple reaction time (SRT) tasks to compute the decision duration (Carsten et al., 2023; Cisek et al., 2009; Derosiere et al., 2022; Thura, 2020), but as we were using different constraints on movement speed, we chose the CRT in order to obtain a measure of the time required to prepare each type of movement (i.e., for free, fast and slow movements). Indeed, there is some evidence that even if subjects prepare their movement in advance in both the SRT and the CRT tasks, their level of preparation is lower in the CRT than in the SRT (Quoilin et al., 2019). Then, *decision accuracy* was calculated as the percentage of trials in which subjects choose the correct lateral circle, referred to as *%Correct*. Thanks to the UGM, we also obtained the intercept and the slope of an urgency signal for each participant and each type of block, as an additional primary endpoint measure. These two parameters then enabled us to estimate the evolution of the *urgency level* over time. To characterize the speed at which subjects moved, we considered the time it took for participants to perform the index flexion movements. Hence, *movement duration* corresponds to the mean duration of a single tap averaged over the four flexions participants performed in each trial of the task.

#### Statistical analyses

##### Experiment 1

The purpose of this experiment was to further investigate the existence of a shared invigoration process linking decision and movement by looking at how contextual changes in decision speed influence movement speed.

###### Effect of decision-related instructions on decision endpoint measures

In order to check whether subjects followed instructions and changed their decision policy between fast and slow decision blocks, we first ran paired t-tests on decision duration and decision accuracy. We then considered the urgency signal estimated for each type of block, running a mixed model with BlockType (fast decision, slow decision) as a factor, Time (elapsed during the decision phase) as a covariate and subject number as a cluster variable (*Urgency* ∼ *BlockType* * *Time* + (*BlockType* * *Time*|*Subject*)). Since we hypothesized a relationship between urgency and decision duration that would hold true across subjects, the covariable was not scaled.

###### Effect of decision-related instructions on movement endpoint measures

To investigate the context and time-sensitive effect of decision duration, we ran general mixed models on movement duration with BlockType (fast decison, slow decision) as a factor, decision duration as a covariate and subject number as a cluster variable (*Movement duration* ∼ *BlockType* * *Decision duration* + (*BlockType* * *Decision duration*|*Subject*)). Our hypothesis was that movement and decision duration correlate within-but not necessarily between-subjects. Hence to get rid of individual average differences in decision duration, we used Z-scores clusterwise to scale the covariable (i.e., decision duration). Post-hoc comparisons were conducted using t-tests with Bonferroni correction for multiple comparisons. Monotonic relationships between block-related changes in movement and block-related changes in urgency/decision were tested using Spearman’s rank correlations.

##### Experiment 2

Here, we aimed at investigating the reverse relationship, as compared to Experiment 1; that is, how contextual changes to movement speed influence decision speed. In addition, this experiment aimed at testing the hypothesis that such a link is masked when detrimental to the reward rate.

###### Effect of movement-related instructions on movement endpoint measures

In order to check whether subjects followed instructions in both groups and changed their movement speed between fast and slow movement blocks, we performed a two-way ANOVA_RM_ on movement duration with BlockType (fast movement, slow movement) and Group (fast decision group, slow decision group) as within-subject factors.

###### Effect of movement-related instructions on decision endpoint measures

The analyses for Experiment 2 were similar to those for Experiment 1, except that here, we looked at the reverse relationship by running a general mixed model on decision duration with BlockType (fast movement, slow movement) and Group (fast decision, slow decision) as factors, movement duration as covariable and subject number as a cluster variable (*Decision duration* ∼ *Group * BlockType* * *Movement duration* + (*Group * BlockType* * *Movement duration*|*Subject*)).

### Exploratory investigation of pupil dilation

In Experiment 1, we recorded pupil diameter from 19 of the 62 participants during task performance. However, one participant did not finish the experiment, so pupil data were analyzed on 18 participants (11 Women, 24.5 ± 3.7 years old). This allowed us to investigate the involvement of the arousal system (proxied by pupil size) in generating the urgency signal (McGinley et al., 2015; Wagenmakers et al., 2018). To this end, the pupil diameter was acquired with an EyeLink 1000+ eye tracker video-based system (SR Research), recording monocularly pupil size (in arbitrary units) and eye movements with a sampling frequency of 1000 Hz. Before analyzing the data, pupil traces were preprocessed to remove eye-blinks, which were identified by the blink detection algorithm implemented in Matlab (https://github.com/alexandre-zenon/pupil/), and replaced by linear interpolations. Furthermore, traces were first filtered by a high-pass filter of 0.01 Hz, then downsampled to 10 Hz to facilitate the analysis. In order to characterize block-related changes in pupil size during the decision phase, we extracted the peak of pupil dilation following the first flexion initiation in each trial (see Fig. S3 in supplementary materials). We then analyzed these Peak pupil data by means of a mixed model with BlockType (fast decision, slow decision) as a factor, decision duration as covariable and subject number as a cluster variable (*Peak pupil* ∼ *BlockType* * *Decision duration* + (*BlockType* * *Decision duration*|*Subject*)). Similarly to the urgency signal analysis, we did not scale the covariable. This analysis allowed us to investigate whether Peak pupil size would evolve as a function of decision duration, as shown for urgency, both in the fast and slow decision blocks. Then, in an additional analysis aimed at further characterizing the evolution of Peak pupil size over time elapsed during deliberation, we obtained for each subject the mean Peak pupil dilation in 8 bins of decision duration (from short to long) and fitted these pupil data with a linear regression to obtain an intercept and a slope. These pupil parameters (intercept and slope) were put in relation to those extracted from the urgency signal (intercept and slope) with Spearman’s rank correlations.

## Results

### Experiment 1

#### Subjects adjusted their decision duration according to the block instruction

In this experiment, we had participants perform the Tokens task in fast and slow decision blocks, which required them to favor either decision speed or accuracy, respectively. Consistently, participants took less time to decide in the fast (1647 ± 276 ms) than in the slow decision blocks (2220 ± 111 ms; t_1,55_ = - 15.236, p < 0.001, Cohen’s d = - 2.036, see Fig. 2.B). This shortening in decision duration influenced choice accuracy: participants were less accurate when deciding in the fast decision blocks (84 ± 5 % of correct choices) than in the slow decision ones (94 ± 2 %; t_1,55_= - 14.175, p < 0.001, Cohen’s d = - 1.894). These findings indicate that participants followed the instructions by adopting a strategy favoring speed in the fast decision blocks and accuracy in the slow decision blocks.

Block-related changes in decision duration were reflected in the level of urgency (Thura et al., 2014; Derosiere et al., 2022). Indeed, the mixed model analysis on the urgency signal revealed a main effect of the factor BlockType (see Fig. 4.A, F_1,55_ = 101, p < 0.001): as expected, the level of urgency was on average higher in the fast (0.327 ± 0.166 a.u) than in the slow decision blocks (0.059 ± 0.107 a.u.). Importantly, this effect was also true when considering the decision period in separate temporal bins (see Table S2 in supplementary materials): the urgency level was higher in the fast than in the slow decision blocks, regardless of the bin considered, up to the last temporal one (all t_1,55_ > 3.760, all p < 0.001, all Cohen’s d > 0.500). Moreover, the mixed model analysis revealed a significant effect of the factor Time (F_1,55_ = 112, p < 0.001), indicating that urgency increased as the time elapsed during deliberation in both fast and slow decision blocks, as also previously shown (Cisek et al., 2009; Thura et al., 2014; Derosiere et al., 2022; Murphy et al., 2016; Steinemann et al., 2018). Finally, we observed a significant BlockType x Time interaction (F_1,951_ = 475, p < 0.001), suggesting that the time course of urgency differed between the two block types. To further address this point, we considered the intercept and the slope of the urgency signal (estimated with the urgency gating model) in both block types. Interestingly, paired-t-tests revealed a significantly higher intercept in fast decision blocks (0.195 ± 0.140 a.u.) compared to the slow decision ones (- 0.200 ± 0.251 a.u.; t_1,55_ = 11.097, p < 0.001, Cohen’s d = 1.483), indicating a greater urgency at decision onset in the former blocks. However, the urgency exhibited a slower increase in the fast than in the slow decision blocks. That is, although the slope was significantly positive in both block types (both t_1,55_ > 5.000, both p < 0.001, both Cohen’s d > 0.700), it was nevertheless smaller in the fast (0.097 ± 0.125 a.u.) than in the slow decision blocks (0.192 ± 0.119 a.u.; t_1,55_ = - 5.241, p < 0.001, Cohen’s d = - 0.700). Hence, the urgency was generally higher when subjects were in a context promoting fast decisions, with a peak difference at decision onset; from there on urgency increased over time in both block types but did so slower in the fast decision blocks, thus reducing the gap with the slow decision blocks, although the difference remained significant until the end of the decision process.

**Figure 4.**
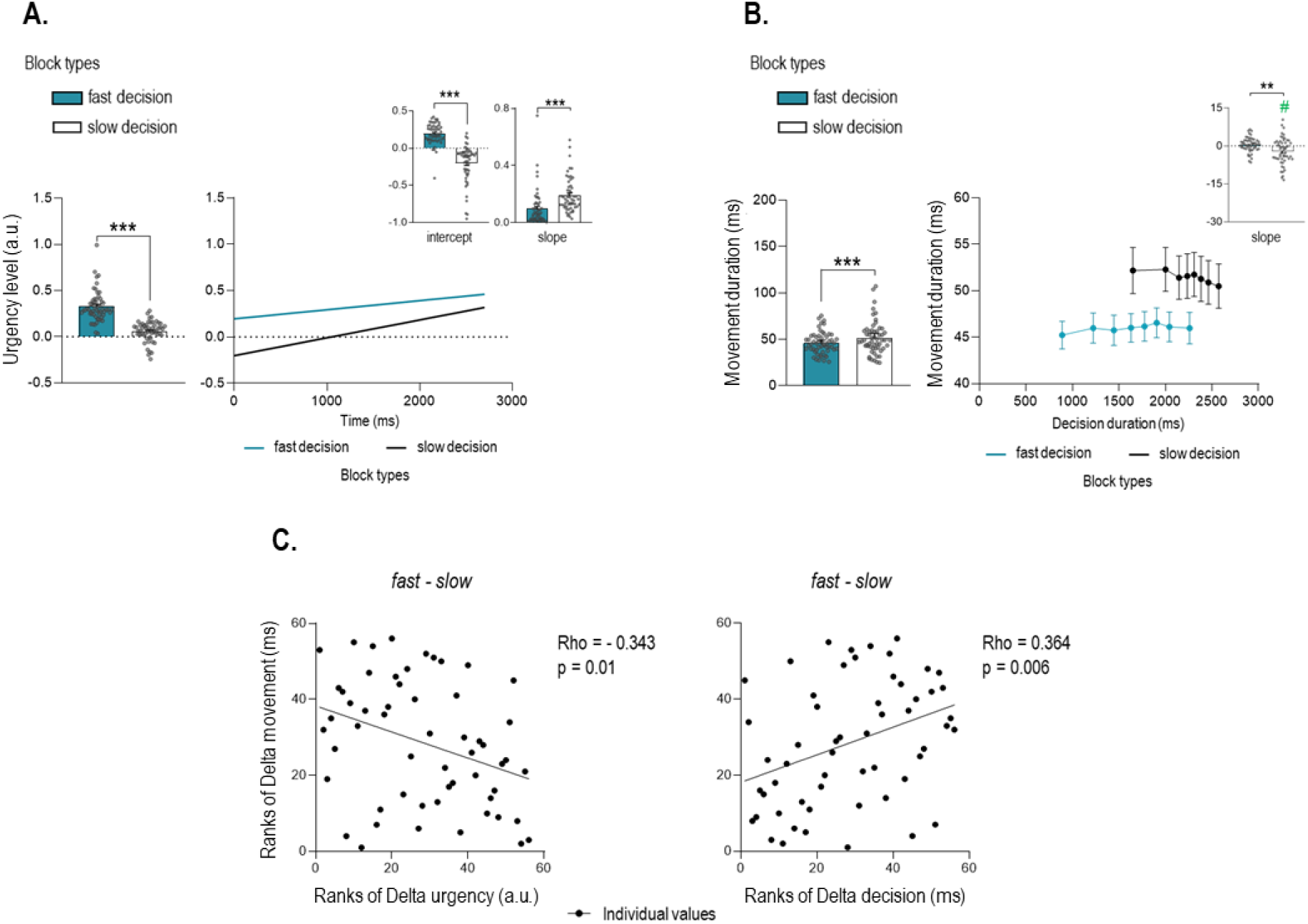
Results for Experiment 1. **A. Urgency signal.** The block-related changes in decision duration were reflected in the level of urgency (left panel), with a higher level of urgency in fast (blue bar) than slow decision blocks (white bar). Consistent with the literature, the evolution of the urgency signal over time (right panel) depended on the type of block: it was higher at decision phase onset (greater intercept, top left inset) and increased more slowly (lower slope, top right inset) in fast compared to the slow decision blocks. **B. Movement duration.** Decisions were implemented by shorter (faster) movements in fast decision compared to slow decision blocks (left panel) even if such a coregulation was not beneficial to the reward rate. Notable, we also found that movement duration shortened with increasing decision duration, but this was only true in slow decision blocks (negative slope, the inset) **C. Correlations.** The more participants responded with a high level of urgency in fast decision compared to slow decision blocks, the more their movements became shorter (faster) in the fast compared to the slow decision blocks (left panel). Similarly, those participants with the largest reduction in decision duration were also those who shortened their movements the most in the fast compared to the slow decision blocks (right panel). Error bars represent SE. # t-test against 0 p < 0.025. ***p < 0.001.

#### Movement duration was influenced by increasing decision urgency

In this first experiment, we assessed the coregulation hypothesis by investigating the impact of changes in decision urgency on movement duration. Based on this hypothesis and in line with prior work (Carsten et al., 2023; Thura et al., 2014; Thura, 2020), we expected that the higher urgency in the fast decision blocks would also fasten the motor response, resulting in shorter (faster) movements in fast decision blocks. Consistently, the mixed model analysis on movement duration revealed a main effect of the factor BlockType (F_1,55.1_ = 24.46, p < 0.001), with shorter values in fast (46 ± 12 ms) than in slow decision blocks (51 ± 18 ms, see Fig. 4.B). Moreover, the analysis revealed a significant BlockType x decision duration interaction (F_1,47.2_ = 11.14, p = 0.002). In order to consider further this interaction, we grouped the movement duration data in 8 bins of decision duration (see Fig. 4.B). Then, we calculated a delta on the global movement duration of the first and the last bins in order to express the change in movement duration (Delta movement). The same delta was calculated on decision duration (Delta decision) to finally compute a slope representing how much movement duration varied as a function of changes in decision duration (i.e., Delta movement /Delta decision). A negative slope would indicate a shortening of movement duration with increasing decision duration while a positive slope would reflect the opposite relationship, that is, a lengthening of movement duration with decision duration. Interestingly, the slope was negative in the slow decision blocks (- 2.11 ± 5.14; t_1,55_ = - 3.076, p < 0.025 with Bonferroni correction, Cohen’s d = - 0.411 when compared to 0 using Student’s t-tests) but it was null in the fast decision blocks (0.438 ± 2.779; t_1,55_ = 1.180, p = 0.243, Cohen’s d = 0.158). Consistently, the slope was found significantly lower in the slow than in the fast decision blocks (paired t_1,55_ = 3.159, p < 0.01, Cohen’s d = 0.422). Hence, altogether the analyses indicate globally shorter movement durations in the fast (higher urgency) than in the slow decision blocks but with an acceleration of movement execution over deliberation time occurring in the latter but not in the former decision blocks.

We then continued to investigate the relationship between decision urgency and movement duration by means of correlations. To do so, we computed deltas of movement duration between the fast and slow decision blocks (delta fast-slow decision blocks) and considered the degree to which changes in movement duration between the two block types were related to the instructed changes in decision speed. To characterize the latter, we computed both the delta of urgency and the delta of decision duration, as depicted on the left and right sides of Figure 4.C., respectively. Interestingly, Delta movement duration correlated negatively with Delta urgency (Rho = - 0.343, p = 0.01, 95%-confidence intervals [CI] = [-0.553, - 0.079]), indicating that the more urgency got high in fast compared to slow decision blocks (higher delta values), the more movements became short (fast) in this type of block (lower delta values). Considering deltas in decision duration instead of urgency led to the same result, which here was manifest as a positive correlation (Rho = 0.364, p = 0.006, CI = [0.114, 0.577]): the more subjects shortened their decision duration in fast compared to slow decision blocks (lower deltas), the more the movements became shorter (lower deltas). Altogether, these findings are consistent with a coregulation of decision and movement speed (Carsten et al., 2023; Spieser et al., 2017; Thura, 2020).

### Experiment 2

#### Subjects adjusted their movement duration according to the block instruction

In this experiment, we had participants perform the Tokens task in fast and slow movement blocks, which required them to report their decision with either fast or slow index finger flexions. Consistent with these instructions and as shown on Figure 3.B, the ANOVA_RM_ revealed that movements were on average of shorter duration in the fast (37 ± 11 ms) than in the slow movement blocks (59 ± 24 ms; F_1,53_ = 106.879, p < 0.001, partial eta-squared (ɳ_P_^2^) = 0.673). Moreover, participants in this experiment were split in two groups where they were implicitly encouraged to either favor decision speed (fast decision group) or decision accuracy (slow decision group). Participants in both groups displayed equivalent movement durations according to the specific instructions of the two block types, as indicated by the absence of factor Group or BlockType X Group interaction on movement duration (all F_1,53_ = [0.111, 1.257], p = [0.267, 0.740], ɳ_P_^2^ = [0.002, 0.024]).

#### Decision duration was influenced by movement duration

In this experiment, we assessed the coregulation hypothesis by investigating the impact of block-related changes in movement duration on decision urgency. Based on this hypothesis and in line with prior work (Carsten et al., 2023; Kita, et al.,2023.), we expected that the requirement to move fast would naturally increase the urgency, resulting in shorter (faster) decisions in fast than slow movement blocks. Yet, as explained in the introduction, we expected that this should only occur if such a coregulation is not detrimental to the reward rate. In other words, we predicted that moving faster in the fast movement blocks would only translate into faster decisions (higher urgency) in the group favoring decision speed anyway (fast decision group) but not in the group prioritizing decision accuracy (slow decision group).

Unsurprisingly, the mixed model analysis revealed a significant effect of the factor Group on decision duration (F_1,55.9_ = 6.32, p = 0.015). As such, decision durations were shorter in the fast decision group (1997 ± 175 ms) than in the slow decision group (2104 ± 158 ms), as implicitly promoted in this experiment. Furthermore, we found a significant Group x BlockType interaction on decision duration (F_1,143.3_ = 6.82, p = 0.010; see Fig. 5.A). As such, changes in movement duration from the slow to the fast movement blocks elicited distinct effects on decision duration depending on the group. That is, post-hoc analyses revealed that participants in the fast decision group made shorter decisions in fast (1990 ± 183 ms) than slow movement blocks (2004 ± 170 ms; t_161.1_ = 2.957, p < 0.025 with Bonferroni correction). Such an effect was not found in the slow decision group, where decision durations were not significantly different between the fast (2112 ± 164 ms) and slow movement blocks (2096 ± 154 ms; t_126.4_ = - 0.667, p = 1.000). Hence, faster movements led participants to decide faster when such a coregulation was consistent with their task goal, in the fast decision group, but not when it would have been detrimental, in the slow decision group.

**Figure 5.**
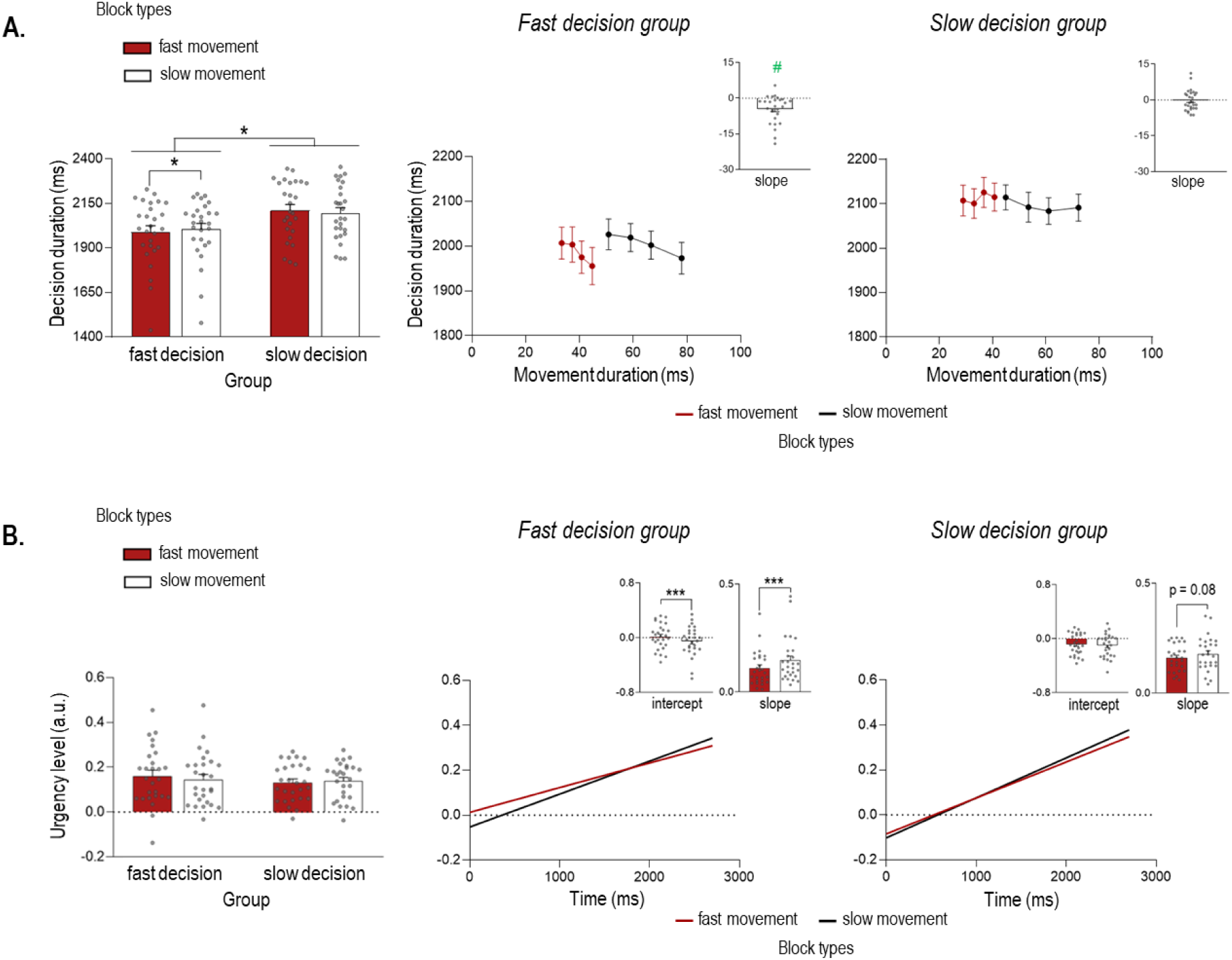
Results for Experiment 2. **A. Decision duration.** As expected, participants in the fast decision group made faster decisions than those in the slow decision group (left panel). Moreover, as also shown on the left panel, only in the fast decision group were decision durations affected by the movement block type, with shorter (faster) decisions accompanying faster movements (red bar). On the right panel you can also see that long movement durations were associated with short decision durations only in fast decision group (see insets displaying specifically the slope in the fast and slow decision groups). **B. Urgency signal.** The overall urgency level did not differ significantly between the two groups and the two types of block (left panel). However, as shown on the right panel, its evolution over time differed between the two groups of subjects. More precisely, the movement block type affected the intercept and the slope of the urgency signal in the fast decision group with only a marginal effect on the slope in the slow decision group. Indeed, the urgency signal was higher at the decision phase onset (greater intercept, top left inset) and increased more slowly (lower slope, top right inset) in fast (red bar) block relative to slow (black bar) movement blocks in the former group only. Error bars represent SE. # t-test against 0 p < 0.05, *p < 0.05, ***p < 0.001.

The mixed model analysis also revealed a main effect of the factor movement duration (F_1,16459.4_ = 8.21, p = 0.004) on decision duration. Similar to the analysis run in Experiment 1, we grouped decision duration in bins of movement duration (here we used 4 bins) in order to calculate a delta between the first and the last bins on decision (Delta decision) and movement durations (Delta movement). These deltas were then used to compute a slope expressing changes in decision duration as a function of changes in movement duration (Delta decision/Delta movement). A Student’s t-test against 0 showed that the slope was significantly negative (t_1,53_ = - 3.152, p < 0.01, Cohen’s d = - 0.429) indicating that long movement durations were associated with short decision durations. However, this effect interacted with the factor Group (Movement duration x Group F_1,16459.4_ = 5.00, p < 0.05). More precisely, following Student’s t-tests against 0, the slope was significantly negative in the fast decision group (– 4.606 ± 5.824; t_1,26_ = -4.109, p < 0.001, Cohen’s d = - 0.791) but not in the slow decision group (– 0.177 ± 4.392; t_1,26_ = - 0.210, p = 0.835, Cohen’s d = - 0.040). This suggests that, only in the fast decision group, movement duration varied as a function of deliberation time, as observed in Experiment 1. All other effects of the mixed model were nonsignificant (p = [0.092 0.509]).

We then assessed the impact of varying the movement speed in the two different block types on decision urgency itself (see Fig. 5.B). The mixed model analysis revealed a main effect of the factor Time on the urgency signal, consistent with an increasing urgency over time (F_1,47.9_ = 197.94, p < 0.001). Moreover, we found that this main effect interacted with the factor BlockType, indicating that the urgency signal generally increased faster in slow relative to fast movement blocks (BlockType x Time interaction; F_1,916_ = 266.65, p < 0.001). Most interestingly, this interaction depended on the Group (Group x BlockType x Time interaction; F_1,916_ = 32.30, p < 0.001). To better understand this triple interaction, we ran further analyses on the intercept and the slope of the urgency signal (as in Experiment 1). More precisely, paired t-tests revealed a higher urgency intercept in fast (0.013 ± 0.186 a.u.) than slow movement blocks (- 0.053 ± 0.215 a.u.; t_1,26_ = 2.854, p < 0.01, Cohen’s d = 0.549) for the fast decision group, whereas the urgency intercept was around 0.093 ± 0.163 a.u. in both block types for the slow decision group (t_1,26_ = 0.915, p = 0.369, Cohen’s d = 0.176). Hence, urgency was greater at the decision onset in the fast than in the slow movement blocks only for the fast decision group. Yet, it seems that urgency then increased more slowly in fast movement blocks, especially in fast decision group. Indeed, for this group, the urgency slope was lower in fast (0.110 ± 0.079; t_1,26_ = - 3.645) than slow movement blocks (0.147 ± 0.103 a.u.; t_1,26_ = - 3.645, p = 0.001, Cohen’s d = - 0.701). The difference between fast (0.160 ± 0.061 a.u.) and slow movement blocks (0.178 ± 0.078 a.u.) was still present in the slow decision group, although smaller and marginal (t_1,26_ = - 1.83, p = 0.079, Cohen’s d = - 0.351). In other words, the urgency signal was higher at the decision phase onset (greater intercept) and increased slower (lower slope) in fast relative to slow movement blocks in the group pushed to decide quickly while it remained mostly unchanged between the two block types in the group pushed to decide slowly. All other effects of the mixed model were nonsignificant (p = [0.154 0.558]). These results suggest that the instruction to boost movement speed in the fast movement blocks induced a parallel increase of urgency when this was consistent with the task goals, in the fast decision group. By contrast, boosting movement speed did not have such an effect in the slow decision group, when it would have been detrimental. In this case, the subjects decided at a similar speed during both slow and fast movement blocks.

Finally, we investigated how the switch in movement speed between fast and slow movement blocks impacted urgency and decision duration. To do so, we computed, as in the Experiment 1, the deltas between the fast and slow movement blocks (delta fast-slow movement blocks) and looked at how Delta urgency and Delta decision duration correlated with Delta movement duration in both decision groups, as depicted on Figure 6. Interestingly, their relationship differed from the Experiment 1. In the fast decision group, there was no relationship with Delta movement duration whether we considered the Delta urgency (Rho = 0.217, p = 0.276, CI = [- 0.191, 0.538]) or the Delta decision duration (Rho = -0.105, p = 0.601, CI = [- 0.451, 0.318]). In the slow decision group, we also found no significant correlation between Delta urgency and Delta movement duration (Rho = 0.176, p = 0.377, CI = [- 0.213, 0.539]). Surprisingly, we found in this group a relationship between changes in movement duration and decision duration, but here Delta decision duration correlated negatively with Delta movement duration (Rho = -0.510, p = 0.007, CI = [-0.779, - 0.141]), suggesting that participants with shorter movements in fast relative to slow movement blocks (lower deltas) were those who waited the longest to make their decisions (higher deltas).

**Figure 6.**
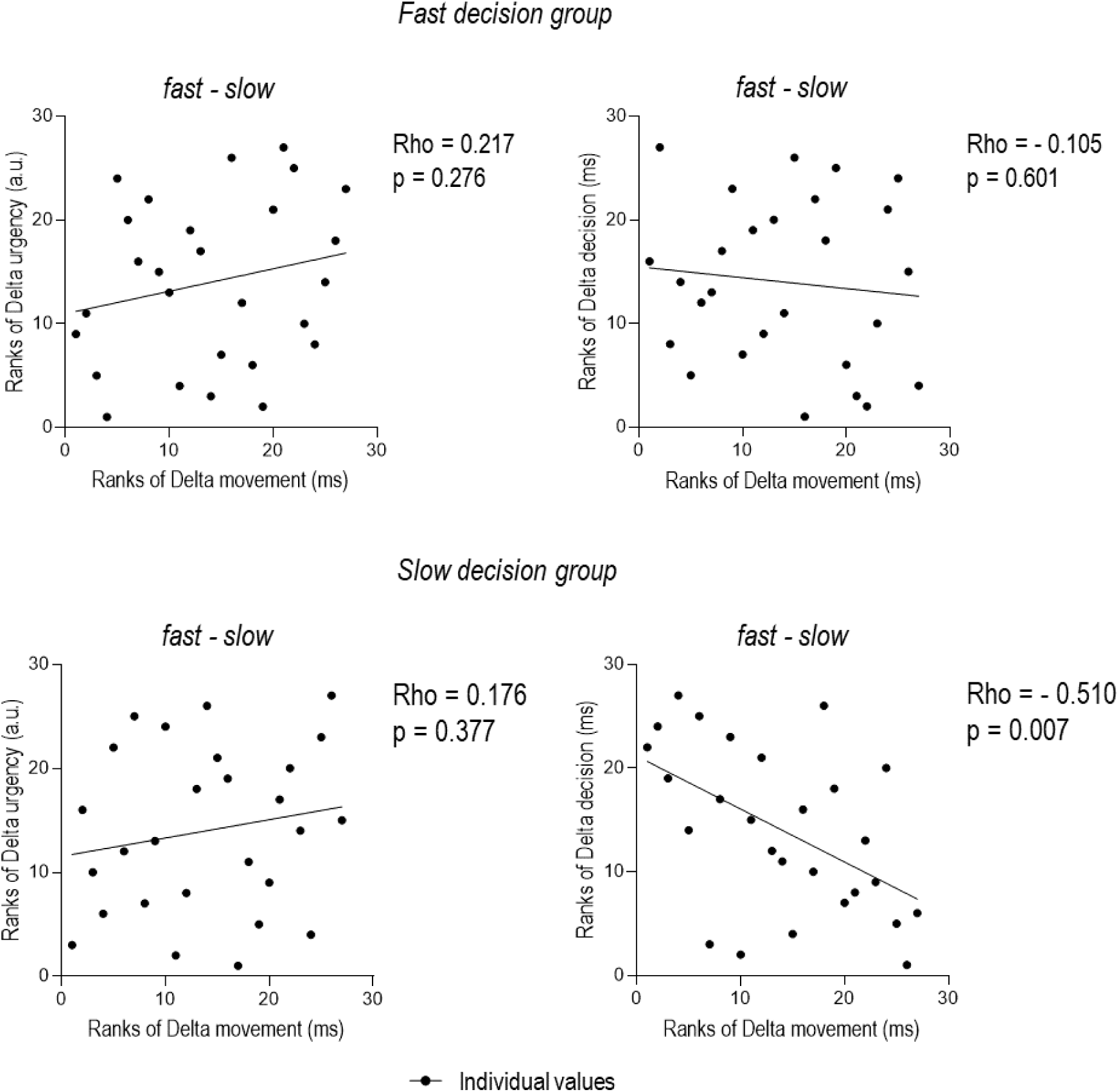
Correlations for Experiment 2. For the fast decision group (upper panel), we did not find any relationship between block-related changes in movement duration (Ranks of Delta movement) and block-related changes in decision features, whether we considered changes in urgency level (Ranks of Delta urgency, left) or changes in decision duration (Ranks of Delta decision, right). For the slow decision group (lower panel), Delta Movement did not correlate with Delta urgency (left) but correlated negatively with Delta decision (right).

In summary, our data in Experiment 2 also indicate a natural coregulation between decision and movement speed when considering the fast decision group in Experiment 2. In contrast, movement speed instructions in the slow decision group did not seem to have any impact on decision speed or on urgency. If anything, the correlations revealed a negative relationship between block-related changes in movement and decision speed in that group favoring accuracy in Experiment 2.

### Analyses of pupil dilation

#### Pupil dilation increased with both urgency and accuracy requirements

Previous studies have shown that increases in urgency induce pupillary dilation (Gross & Dobbins, 2021; Murphy et al., 2016; Reppert et al., 2023; Steinemann et al., 2018). However, the link between pupil dilation and decision urgency is still much debated in the literature. That is, while some have suggested that pupil dilation reflects the involvement of the arousal system in the generation of the urgency signal (Murphy et al., 2016), others have proposed that the urgency-related pupillary dilation rather results from an increased recruitment of the arousal system to ensure decision accuracy under time pressure (Steinemann et al., 2018). In order to directly address this question in the current study, we analyzed pupil dilation as a function of decision duration, in 18 of the subjects who took part in Experiment 1.

The mixed model on Peak pupil dilation did not reveal any significant effect of the factor BlockType (F_1,8257_ = 0.616, p = 0.433; 111 ± 57 a.u. for fast decision blocks and 163 ± 74 a.u. for slow decision blocks; see Fig. 7). However, the analysis revealed a main effect of the factor decision duration (F_1,8265_ = 30.173, p < 0.001), indicating that pupil dilation increased with decision duration in both types of blocks. Interestingly, this effect interacted with the factor BlockType (BlockType x decision duration interaction; F_1,8265_ = 30.173, p < 0.001). In order to understand this interaction, we estimated the intercept and slope of the pupil function in each block type by means of linear regressions (see related section in Methods for more details) and compared them across block types. The intercept, which thus reflects pupil size at the onset of the decision phase did not differ between block types (t_1,17_ = 1.049, p = 0.309, Cohen’s d = 0.247; 85 ± 71 a.u. and 50 ± 152 a.u. in fast and slow decision blocks). However, the slope (reflecting the speed at which the pupil expanded over decision duration) was steeper in slow (50 ± 152 a.u.) relative to fast decision blocks (5 ± 71; t_1,17_ = - 2.107, p = 0.05, Cohen’s d = - 0.497). This result was supported by multiple paired t-tests (Bonferroni corrected) looking at the difference in Peak pupil dilation between fast and slow decision blocks in the 8 bins of decision duration. As such, this additional analysis showed that, in early bins, pupil dilation did not differ significantly between block types but that it became progressively larger in slow than fast decision blocks in the last bins of decision duration (see Table S3 in supplementary material).

**Figure 7.**
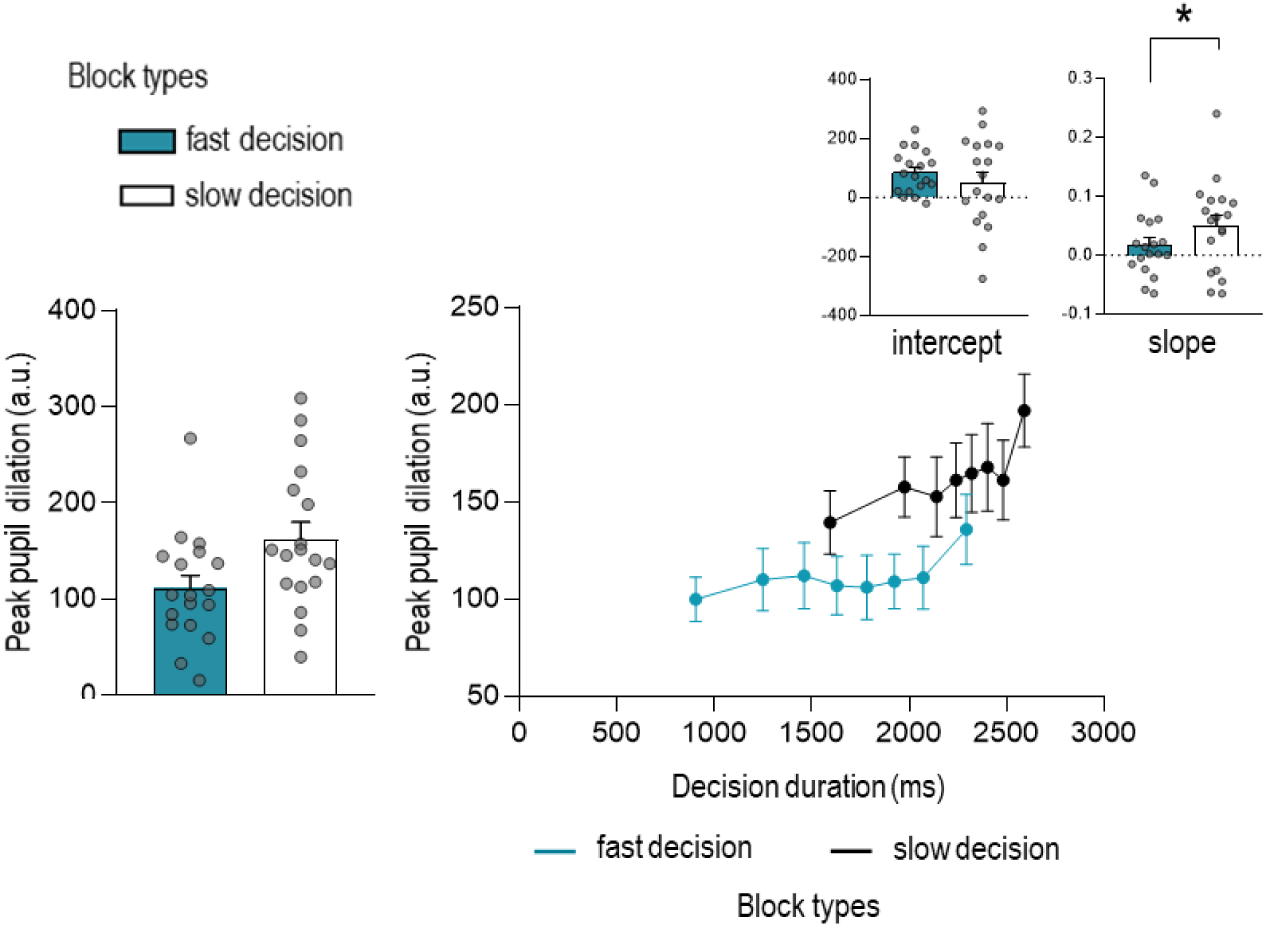
Peak pupil data in Experiment 1 (n=18). Peak pupil dilation was comparable in fast (blue bar) and slow (white bar) decision blocks (left panel). Consistent with the literature, Peak pupil dilation increased with decision duration in both types of blocks (right panel). However, while the pupil dilation seemed similar between the two types of block at the beginning of the decision phase (similar pupil intercepts, top left inset), its dilation tended to increase faster in slow decision blocks relative to fast decision ones (higher pupil slope, top right inset), leading to a progressively more dilated pupil in the former. Error bars represent SE.

To further investigate the link between pupil dilation and urgency, we put into relation the block-related changes in parameters of both functions. In other words, we looked at how block-related changes in pupil intercept (slope) were linked to block-related changes in urgency intercept (slope). These changes were quantified by obtaining delta values (delta fast-slow decision blocks) for the intercept and slope of both the pupil (Delta pupil intercept and Delta pupil slope) and urgency (Delta urgency intercept and Delta urgency slope) functions in all subjects, with thus positive deltas indicating greater values in fast than slow decision blocks. Interestingly, a Spearman correlation on Delta intercept values revealed a tendency toward a positive correlation (Figure 8; Rho = 0.459, p = 0.057, CI = [0.012, 0.779]), suggesting that participants who showed a greater urgency increase at the onset of decisions in fast than slow decision blocks were also those displaying the larger increase in pupil dilation. Moreover, a positive correlation was found when considering the Delta slope values (Rho = 0.478, p = 0.047, CI = [- 0.022, 0.803]), indicating that participants whose urgency signal increased more over time in fast relative to slow decision blocks also showed a larger increase in pupil dilation over time.

**Figure 8.**
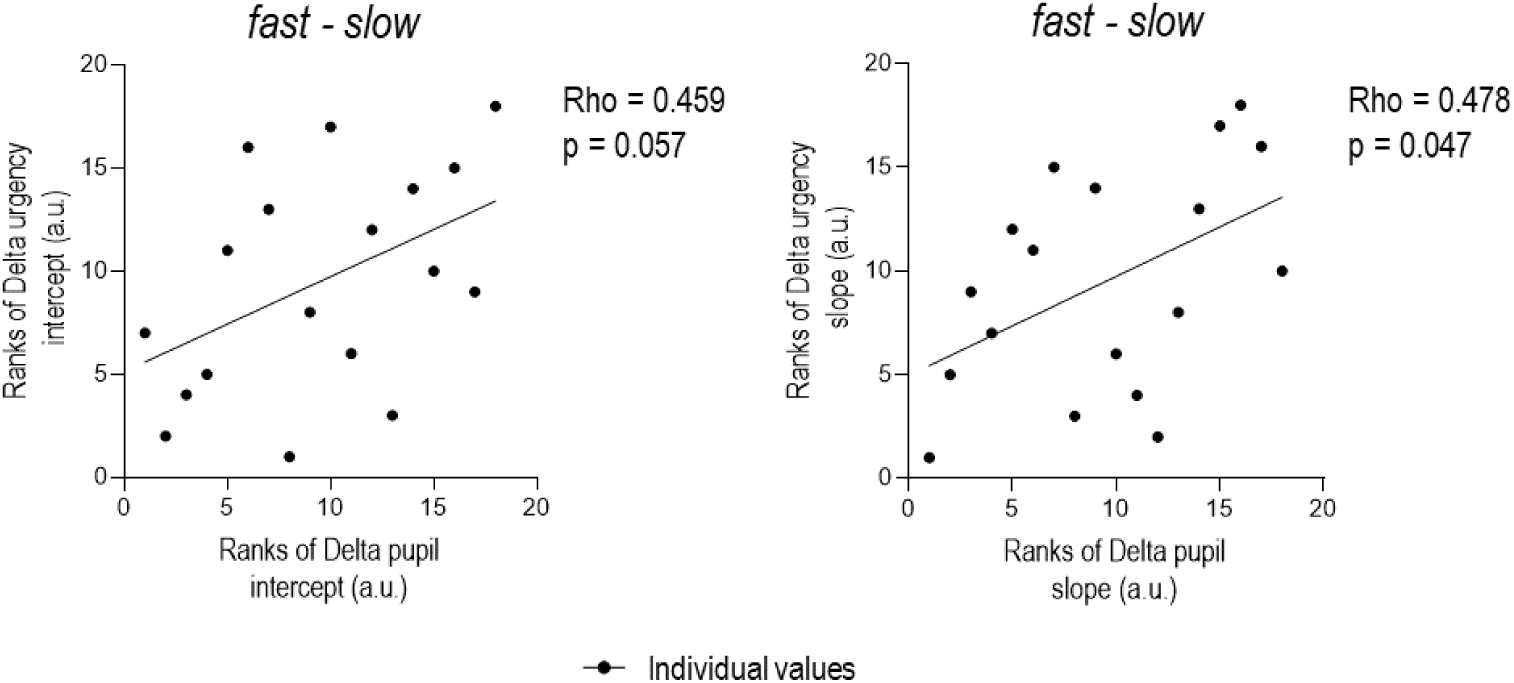
Correlations for Peak pupil data in Experiment 1 (n=18). Block-related changes in the intercept and slope of the Peak Pupil function (Ranks of Delta pupil intercept and slope) were related to block-related changes in the same parameters of the urgency function (Ranks of Delta urgency intercept and slope). That is, the participants showing a greater increase in pupil intercept (slope) in fast compared to slow decision blocks were also those showing a greater increase in urgency intercept (slope).

**Figure 9.**
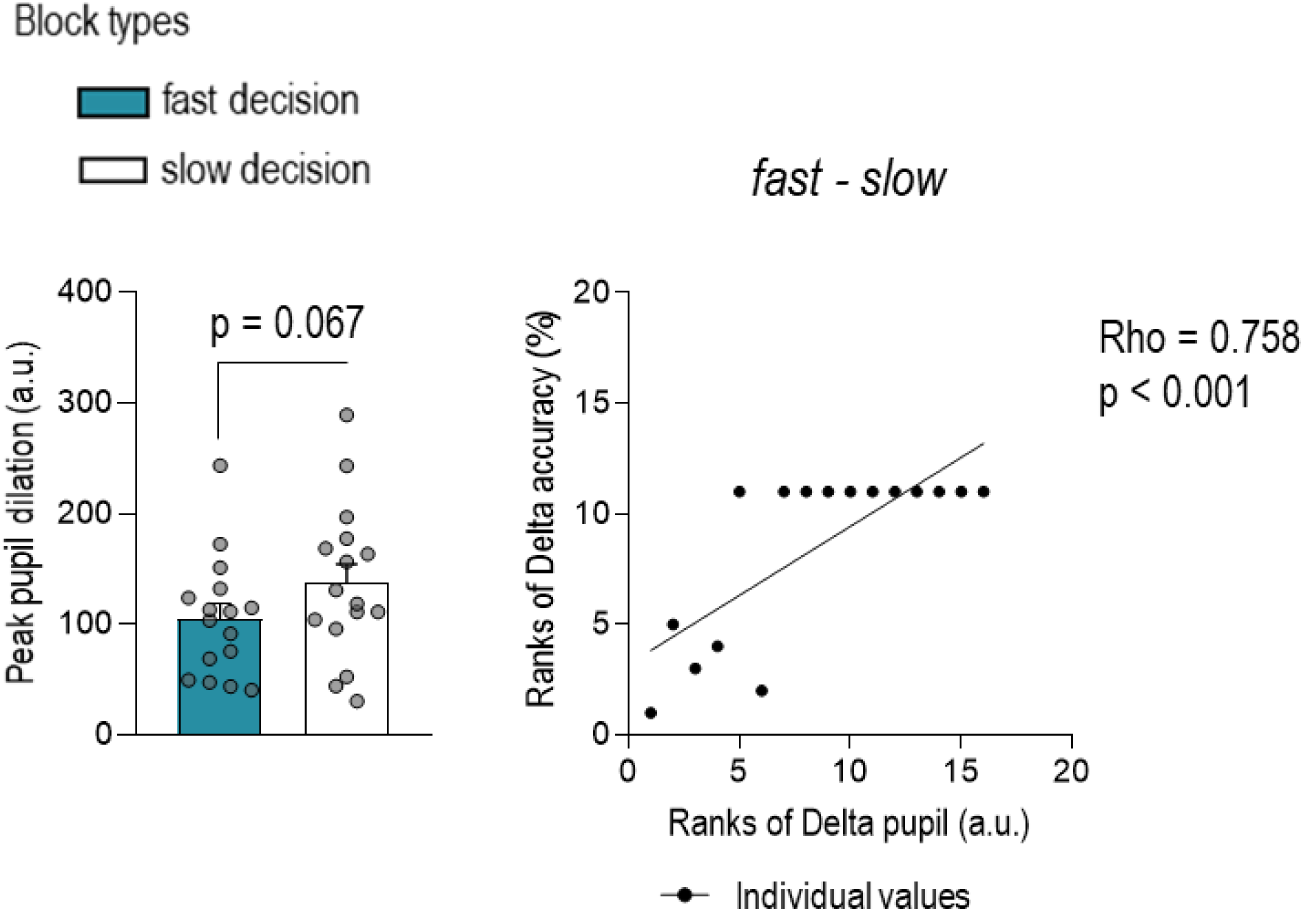
Peak pupil data in ambiguous trials of Experiment 1 (n=16). Peak pupil dilation tended to be larger in slow decision blocks (white bar) than in fast decision blocks (blue bar). Given that deltas were computed by measuring the difference in peak pupil dilation between fast and slow conditions, the more the subjects showed a greater peak pupil in slow decision blocks, the more they displayed a negative delta, which corresponds to a low rank value. Interestingly, we found that the greater this pupil enlargement in slow compared to fast decision blocks (i.e., lower rank value), the greater the gain in accuracy in slow compared to fast decision blocks (i.e., lower rank value).

Altogether, the effects described above are consistent with a tight relationship between urgency and pupil dilation. Indeed, both urgency and pupil dilation clearly increased with decision duration, and more so in slow decision blocks. Moreover, block-related increases in urgency intercept and slopes were associated with parallel enlargements in pupil size (though marginal for the intercept). However, we also found that, pupil size tended to be smaller in fast than in slow decision blocks (and significantly so in later bins), contrary to urgency which showed the reverse effect. Such an effect on the Peak pupil data was not related to differences in baseline (tonic) levels of pupil dilation as these values were equivalent in both block types (see Fig. S4 in supplementary materials). Nor was this effect due to the fact that decision durations differed across block types within the same bins (see Fig. 7). Indeed, when considering only a set of equivalent ambiguous trials with long decision durations (i.e., decisions falling between Jump_-9_ and Jump_-10_) and a constant level of evidence (i.e., with a success of probability at decision time equal to 0.8125 in all trials), we found that pupil dilation remained greater in slow decision blocks, although the difference with fast decision blocks was marginal (137 ± 70 a.u. and 105 ± 54 a.u., respectively; t_1,15_ = 1.97, p = 0.067, Cohen’s d = 0.493; 2 subjects had to be excluded from this analysis, because of a lack of data).

We then considered the possibility that the larger pupil size observed in slow decision blocks was due to a special focus on accuracy in this condition. As such, subjects had a fixed number of trials here (contrary to fast decision blocks) and they then plausibly recruited all available resource’s to be as accurate as possible, including increased arousal. To test this hypothesis, we considered again the subgroup of ambiguous trials described above for which we computed a block-related delta (fast-slow) for the accuracy and pupil data. Interestingly, in those particular trials, the Delta pupil size was positively correlated with the Delta accuracy (Rho = 0.758, p < 0.001, CI = [0.501, 0.873]). In other words, the more the pupil was dilated in the slow decision blocks compared to the fast decision blocks (i.e., the more the delta pupil was negative), the more subjects exhibited better accuracy in the slow than in the fast blocks (i.e., the more the delta accuracy was negative). In conclusion, variations in pupil-related arousal in the current study seemed to be driven by two overlapping factors: the level of urgency, as expected, and the strategic allocation of resources to promote accuracy.

## Discussion

The aim of our study was to provide a principled explanation to the relation between decision and movement speed. Consistent with this, our data supports the hypothesis that coregulation operates as a default mode. However, our findings also show that this form of control can fade or even be replaced by a tradeoff relationship, depending on the task goals. Based on these behavioral observations, we propose that several processes come into play to shape the speed of our decisions and movements either selectively or jointly, the latter mode being best observed when a default coregulation of decision and movement allows to maximize the reward rate, or at least, when it is not detrimental to it.

### The relationship between decision and movement speed adapts to context-dependent task goals

Context-dependent changes in decision speed, as elicited in Experiment 1, led to clear changes in movement speed that are consistent with a coregulation of both levels of control. Indeed, subjects made faster movements in fast decision blocks than in slow decision blocks, and these changes in movement speed were correlated with the strength of changes in decision speed: the more subjects adjusted their decision speed according to the instruction, the more they showed a parallel change in movement speed. Note that, similar to Carsten et al (2023), changes in movement speed accompanied block-related variations in decision speed despite its absence of impact on the reward rate, substantiating the view that such coregulation type of control occurs naturally.

Context-dependent changes in movement speed, as required in Experiment 2, also affected decision speed. Yet, the direction of the effect differed according to the group that was considered. That is, the group whose reward rate was maximized by a strategy favoring decision speed (i.e., the fast decision group) showed faster decisions in fast movement compared to slow movement blocks. Even if this coregulation effect was not as strong as in Experiment 1, it was still significant, pointing to a similar form of coregulation. In contrast, the group whose reward rate was maximized by a strategy favoring decision accuracy (i.e., the slow decision group) did not show such a covariation of decision speed between the slow and fast movement blocks. Quite the contrary, in this group we even observed the reverse relationship. More precisely, although decision speed was similar between fast and slow movement blocks, the changes in movement speed correlated negatively with changes in decisions speed: the more subjects accelerated their movement speed in fast compared to slow movement blocks, the more they slowed down their decision. This is consistent with a tradeoff type of relationship in this (slow decision) group, contrasting with the coregulation effect observed in the other (fast decision) group, despite the fact that subjects from both groups displayed a coregulation effect when manipulating decision speed in Experiment 1.

Interestingly, similar observations regarding context-dependent variations in Experiments 1 and 2 were made when considering directly the relationship between movement speed and decision urgency itself (and not only decision speed), as done for the first time in this study. As such, in Experiment 1, the decision urgency was clearly higher in fast decision blocks compared to slow decision blocks and this upregulation was found to significantly correlate with the changes in movement speed that occurred between the two contexts: the greater the context-dependent increase in urgency level, the more subjects accelerated their movement speed from one block type to the other. Similarly, the coregulation observed in the fast decision group of Experiment 2 was also apparent when considering the urgency: fast movement blocks were associated with a higher urgency intercept than slow movement blocks in this group of individuals. Such a relationship was not found when the other (slow decision) group was considered. Hence, altogether, these findings indicate that the relationship between decision speed/urgency and movement speed can shift from a coregulation to a tradeoff according to the task goals, which provides an explanation for the apparent discrepancy between findings of past studies (Carsten et al., 2023; Kita et al.,2023.; Reynaud et al., 2020; Saleri Lunazzi et al., 2021).

### The deliberation time impacts on movement speed

In our experiment, the level of urgency did not only vary in a context-dependent manner (i.e., between blocks) but also in a time-dependent manner (i.e., over the course of a trial). That is, as shown in many past studies, urgency increased during a trial, pushing the subjects to respond as the trial comes to an end. Consistent with a natural coregulation of decision speed and movement, we observed a time-dependent speeding up of movements over the course of a trial, as already reported in past studies (Thura et al., 2014; Thura, 2020; Thura et al., 2012). Interestingly, this was generally true, whether considering results from Experiment 1 or Experiment 2: overall, movement speed increased as deliberation time (and thus urgency) increased, in a given context.

Yet, an intriguing finding here, is that in Experiment 1, this effect was only observed in the slow decision blocks (generally associated with slow movements) but not in the fast decision blocks (where movements were generally faster due to the context-dependent change in urgency elicited by the task instruction to decide fast). A possible explanation for this is that the slope of the urgency function was smaller in the fast than slow decision blocks of this experiment, as shown in the past. That is, when subjects put the emphasis on decision speed, the intercept is higher but then the urgency increases more slowly as evident by a smaller slope in fast decision blocks (Thura et al., 2014; Thura, 2020). Hence, the weaker increase in urgency over the course of the trial in the latter block type may be responsible for the absence of time-dependent effect on movement speed in that condition, compared to the slow decision blocks where the slope is steeper following a lower intercept. Nevertheless, previous studies have shown an acceleration of movements with the passage of time, irrespective of the slope size of the urgency signal (Thura et al., 2014; Thura, 2020). Another plausible explanation is that the impact of urgency on movement speed is counteracted by a process controlling the energy cost. More precisely, the energy expenditure as a function of movement duration is represented by a concave upward function for many types of movement (Shadmehr et al., 2016; Steudel-Numbers & Wall-Scheffler, 2009; Yoon et al., 2018; Zarrugh et al., 1974). As the speed of movement is slower in the slow decision blocks than in the fast decision blocks, it is possible that accelerating the movement speed decreases energy expenditure in the former and increases it in the latter. Increasing the energetic cost of movement in the fast decision blocks would lead to a decrease in the value of the reward, and thus the rate of reward. Therefore, limiting the impact of the urgency signal depending on the energy cost seems entirely credible. The fact that previous studies did not observe a ceiling effect for the urgency signal could be linked to the type of movement involved in the task. Indeed, in these studies, the task involved reaching movements, whereas our experiment required index finger tapping movements. Unlike the latter, reaching movements are large and involve several joints (at least the three articulations of the arm). It is likely that such multi-joint movements offer a wider range of movement speeds than single-joint (tapping) movements.

#### Pupil dilation reflects an increase in arousal related to urgency and decision accuracy

Converse to previous studies (Gross & Dobbins, 2021; Murphy et al., 2016; Steinemann et al., 2018), the pupil dilation did not differ significantly between fast and slow decision blocks. However, we found an increase in dilation as a function of decision duration, suggesting that pupil dilation may be partly linked to the urgency signal (Gross & Dobbins, 2021; Lawlor et al., 2023; Murphy et al., 2011, 2016; Steinemann et al., 2018). This idea is supported by the similarity found between pupil dilation and urgency signal as well as the positive correlations that were found between their parameters (see Fig. S5 in supplementary material). Yet surprisingly, unlike the urgency signal, pupil size tended to be more dilated in slow compared to fast decision blocks. Contrasting with previous studies (Gross & Dobbins, 2021; Murphy et al., 2016; Steinemann et al., 2018), this result suggests a recruitment of the arousal system as a function of both urgency and accuracy. One possible explanation is that the emphasis on decision accuracy in slow decision blocks was particularly high in our design. This pressure could arise from the fact that using a fixed number of trials limits the possibility of increasing the rate of reward, giving importance to each trial. The positive correlation between pupil dilation and performance in ambiguous trials supports this idea, showing that the more the pupil was dilated in slow decision blocks, the more subjects increased their accuracy in this condition for a same level of evidence.

Altogether, these results suggest that pupil dilatation reflects two overlapping actions. More precisely, the recruitment of the arousal system would, on the one side, increase the gain of sensory information at the decision level with the increase in urgency in the fast decision blocks, and on the other side, increase the attention to enhance accuracy in a context where it is prioritized regardless of urgency, as in our slow decision blocks. The mechanisms through which the arousal system can enhance performance accuracy are still unclear, but may involve enhancing processing of sensory information (Zénon, 2019). This hypothesis is in line with a report by Steinemann and his colleagues, who showed that pupil dilation is associated with better encoding of sensory evidence in the visual cortex. Nevertheless, further studies are needed to investigate the involvement of the arousal system in decision making using causal approaches, using for example techniques such as transcutaneous vagus nerve stimulation (Hilz, 2022).

### Several neural mechanisms are likely to shape the relationship between decision and movement speed

#### Plausible neural source underlying the natural coregulation of decision speed and movement speed

A possible candidate for the coregulation effects observed in our two experiments is the basal ganglia (Chen & Yang, 2021; Dudman & Krakauer, 2016; Kaduk et al., 2023; Desmurget & Turner, 2010). As such, this structure could be at the source of a signal invigorating behavior as a whole to maximize reward rate, as hypothesized in several studies of the last decade (Carsten et al., 2023; Cisek & Thura, 2022; Thura, 2020; Thura et al., 2022; Thura & Cisek, 2017). Reward-sensitivity of the dopaminergic system has been extensively studied in the past (Berridge & Robinson, 2016; Salamone & Correa, 2024; Schultz et al., 2017; Weinstein, 2023) Consistently, dopamine seems strongly implicated in the control of decision speed and movement vigor (Bourgeois et al., 2016; Coddington & Dudman, 2019; Niv, 2007; Pietro Mazzoni et al., 2007). Interestingly, it is believed that it amplifies motor gain (Park et al., 2020; Yttri & Dudman, 2018), which seems to rely, at least in part, on inhibitory influences increasing the signal-to-noise ratio in motor neural activity (Duque et al., 2017; Greenhouse, 2022; Vassiliadis et al., 2020; Wilhelm et al., 2022). This dopamine-mediated invigoration may be thus responsible for the increased surround inhibition, a phenomenon observed in studies applying transcranial magnetic stimulation (TMS) during movement preparation, which has traditionally been associated with an increase in movement vigor (Dudman & Krakauer, 2016; Shadmehr et al., 2019) but which has also recently been linked to an increase in decision urgency (Derosiere et al., 2022).

#### Neural source underlying the occurrence of a tradeoff between decision urgency and movement speed

The context-dependent modulations of decision speed as a function of the movement blocks in the slow decision group of Experiment 2 reflects a tradeoff rather than a coregulation of decision and movement speed, despite the fact that there is evidence for a coregulation in this same group in Experiment 1 (see Fig. S6 in supplementary materials). This suggests that the requirement to focus on decision accuracy led this group of subjects to recruit processes allowing to maintain the appropriate level of accuracy despite the fact that switching from slow to fast movement blocks let to a joint invigoration of decision with movement speed. Such hypothesis is supported by a negative correlation found between the UPPS score for urgency and the delta of decision duration in the slow decision group (see Fig. S.7 in supplementary material). Although marginally nonsignificant (p-tendency of 0.083), this correlation suggests that the more impulsive the subjects (higher UPPS urgency score), the less they were able to slow down their decisions (lower delta) in fast movement blocks. In other words, this suggests lesser ability to counteract the effect of common invigoration. Therefore, it is plausible that a cognitive control process (less efficient in impulsive individuals) can come into play and selectively shape decision parameters in a goal-directed manner, helping to “hold your horses”. This type of inhibitory control mechanism has been widely studied during conflict resolution and during action stopping as an extreme case of braking (Frank et al., 2007; Mosher et al., 2021). Several lines of evidence indicate that it relies on the hyperdirect pathway from medial frontal cortex to subthalamic nucleus (Cavanagh et al., 2011; Forstmann et al., 2008; Herz et al., 2017), producing broad, non-specific inhibition in motor areas to slow down the action selection process (Aron et al., 2016; Duque et al., 2016; Klein et al., 2014; Wessel & Aron, 2017). This effect would be associated with a broad motor cortex suppression as already observed in past TMS studies, showing for instance reduced excitability in leg muscles during cautious finger responses (Derosiere et al., 2022; Duque et al., 2017).

## Conclusion

In conclusion, the interaction between decision and movement speed seems influenced by different mechanisms. Our findings highlight two of them; a common invigoration mechanism responsible for a natural coregulation between decision and movement speed, and a cognitive mechanism, which slows down decisions when the reward is linked to the precision of the choices. Contrary to expectations, although the urgency signal was influenced by changes in movement speed, it did not always correlate with it, suggesting that this signal reflects a combination of the different mechanisms involved in decision making and not just that of invigoration. Interestingly, our results support a link between pupil dilation and decision urgency, but here also with the additional influence of an accuracy-promoting factor.

## Acknowledgements

The authors wish to thank Julien Lambert, Benvenuto Jacob, Christian Lebègue, Caroline Quoilin and Justine Hupin for their contribution to the development of the task design. We thank also David Thura for his help on data modeling in the Tokens Task, as well as Claudia Danse, Eline Volders, Baptiste Waltzing and Fanny Schannes for their help with data acquisition.

## Author contributions

FF designed the study, acquired and analyzed the data, and wrote the first draft of the manuscript. IC contributed to the study design and data analyses, TC and GD contributed to the study design, AZ contributed to the data analyses, and JD contributed to the study design and data analyses. All authors contributed to the article and approved the submitted version.

## Supplementary materials

**Figure S1.**
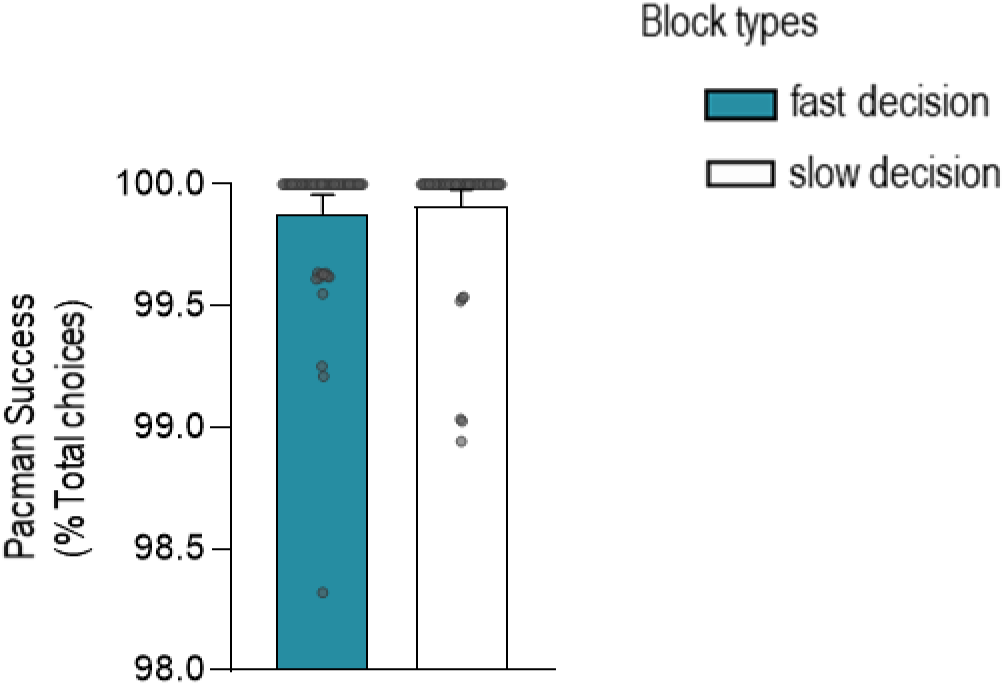
Movement performance in Experiment 1. Participants successfully escaped the ghost (Pacman Success) equally in fast decision blocks (blue bar) and slow decision blocks (white bar). Error bars represent SE.

**Figure S2.**
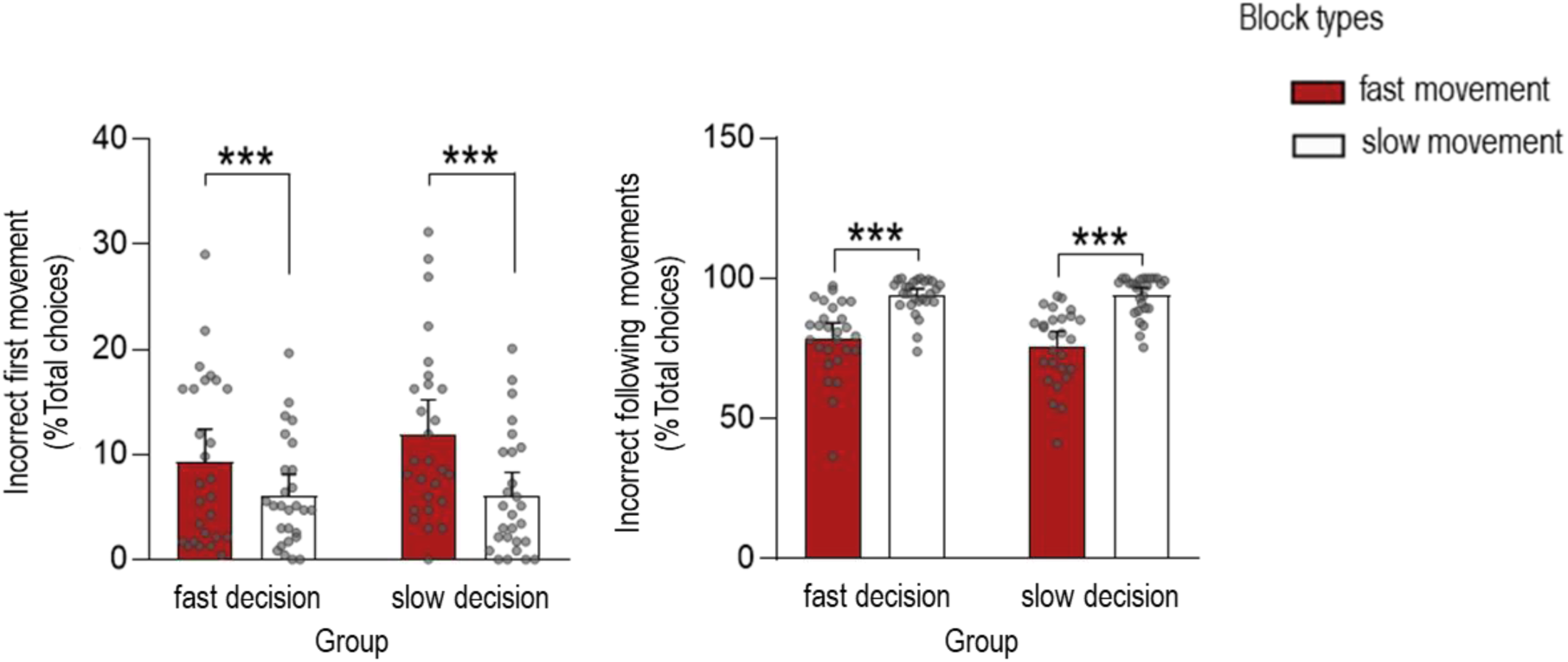
Movement performance in Experiment 2. On average, participants had more difficulty keeping up with the speed imposed in the fast movement blocks (red bars) than in the slow movement blocks (white bars). More precisely, the success rate was lower in fast relative to slow movement blocks for both the first tapping movement (higher percentage of incorrect first movement, left panel) and the following ones in the movement phase (lower percentage of incorrect following movements, right panel). Error bars represent SE. ***p < 0.001.

**Figure S3.**
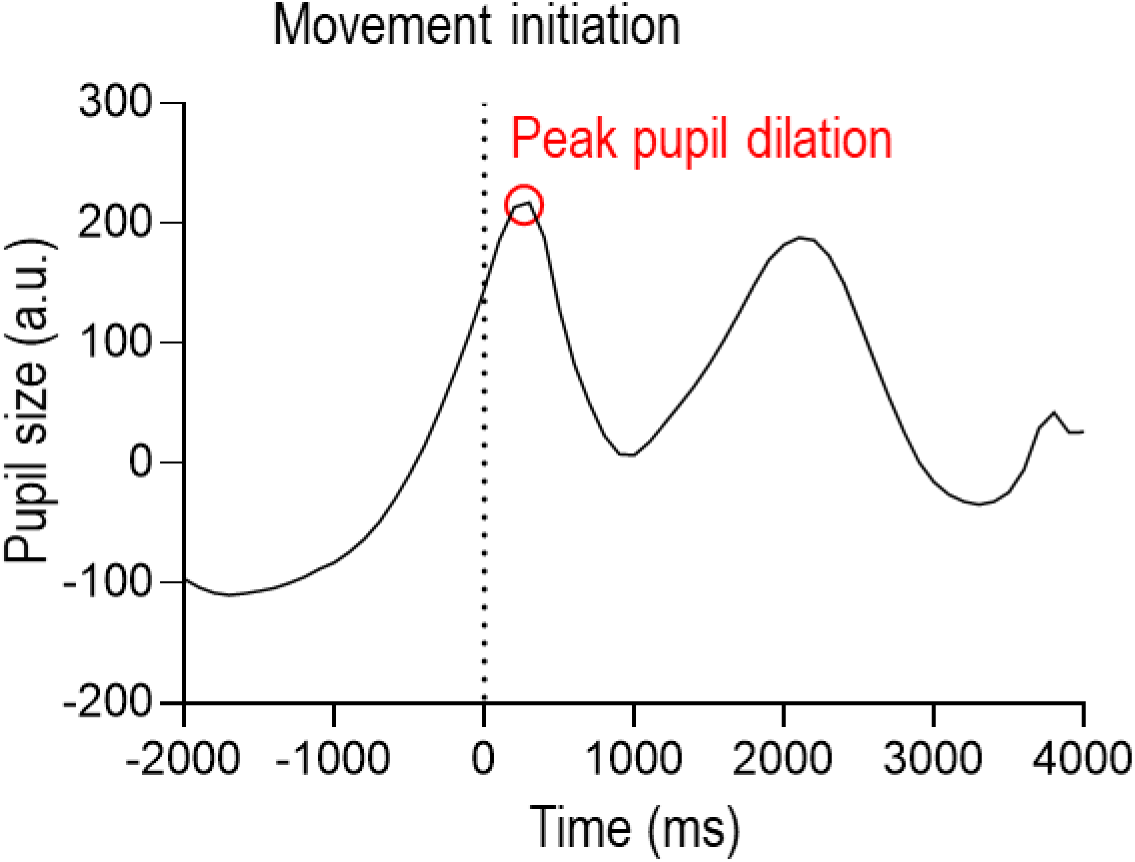
Typical pupil dilation signal. In order to characterize block-related changes in pupil size during the decision phase, we extracted the peak of pupil dilation (Peak pupil dilation) following the first flexion initiation (Movement initiation) in each trial. This figure shows the evoked pupil dilation aligned to movement initiation (0) in one subject.

**Figure S4.**
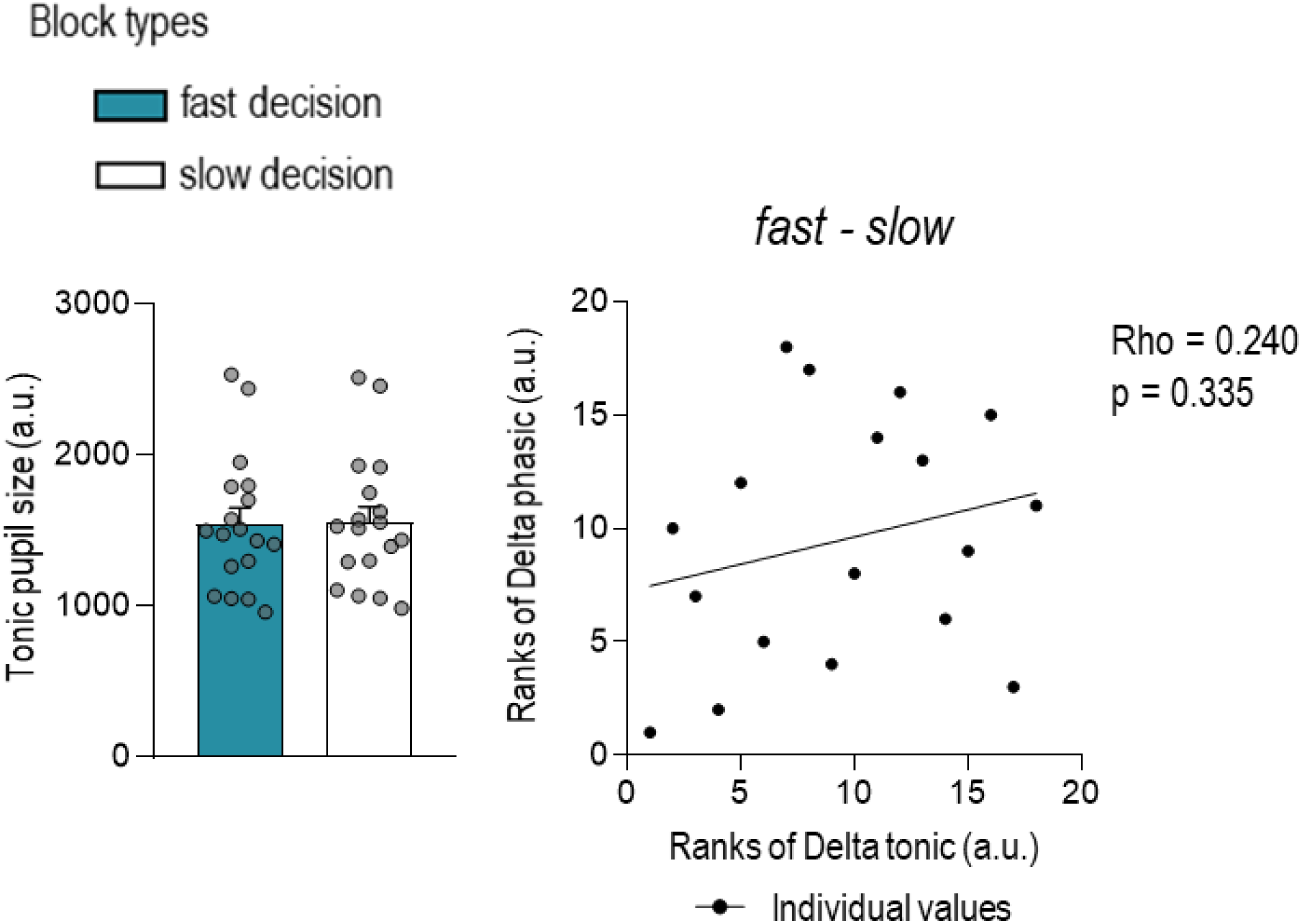
Control analyses on pupil dilation in Experiment 1 (n=18). As a control analysis, we considered a window of 250 ms before the trial onset to measure tonic pupil dilation. As shown on the left panel, such tonic pupil dilation was similar in fast (blue bar) and slow decision blocks (white bar). Furthermore, block-related changes in tonic dilation (Ranks of Delta tonic) did not correlate significantly with change in Peak pupil dilation (Ranks of Delta phasic), ruling out any pupil bias in our results (right panel).

**Figure S5.**
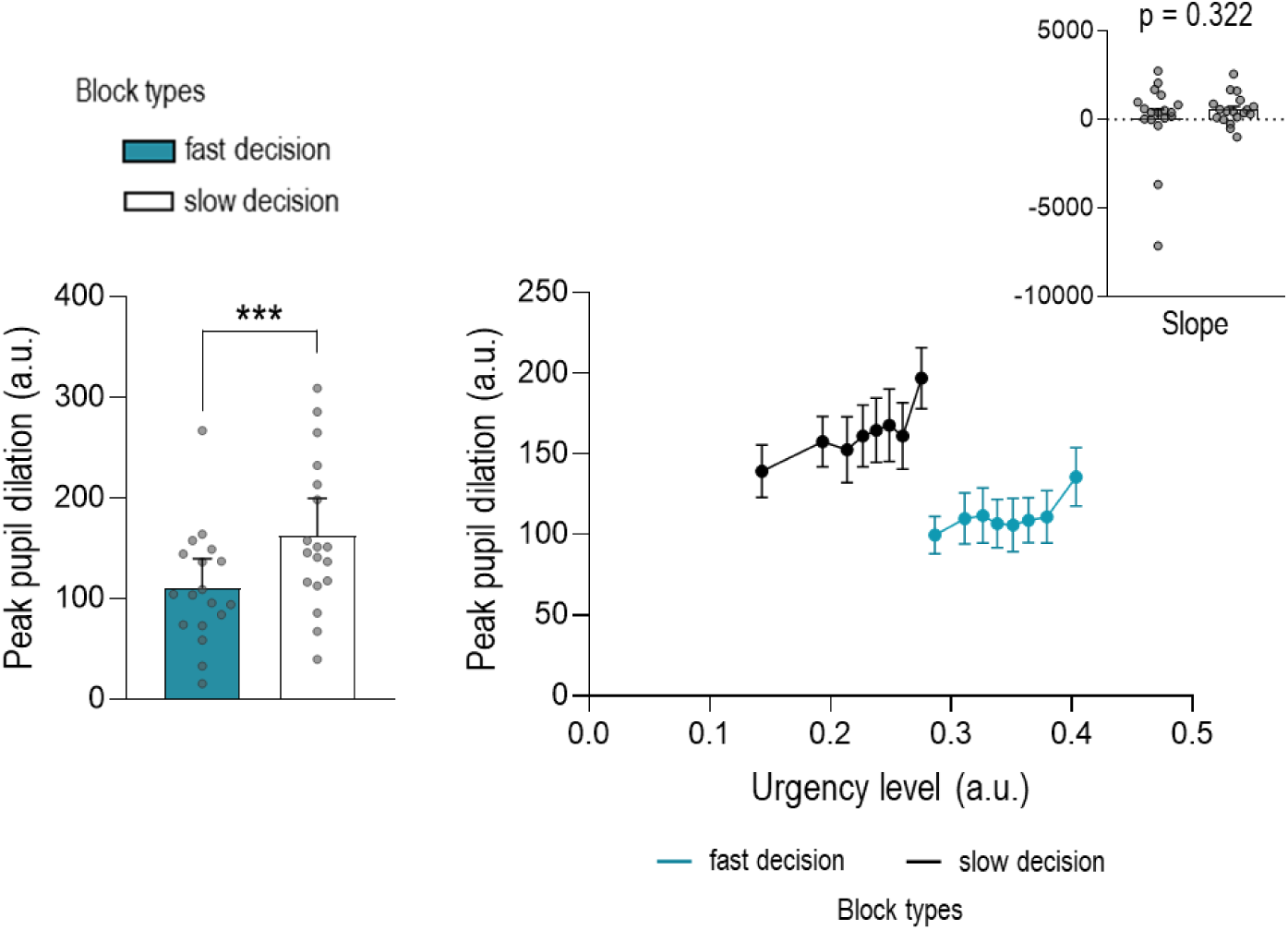
Peak pupil data in Experiment 1 (n=18) as a function of urgency at decision time. Peak pupil dilation was smaller in fast (blue bar) than slow (white bar) decision blocks (left panel). Consistent with the positive correlation found between pupil dilation and urgency signal, Peak pupil dilation increased with urgency level in both types of blocks (right panel). Furthermore, this increase in pupil dilatation with urgency was similar between the two block types as emphasize by the similar slopes (see inset). Error bars represent SE. ***p < 0.001.

**Figure S6.**
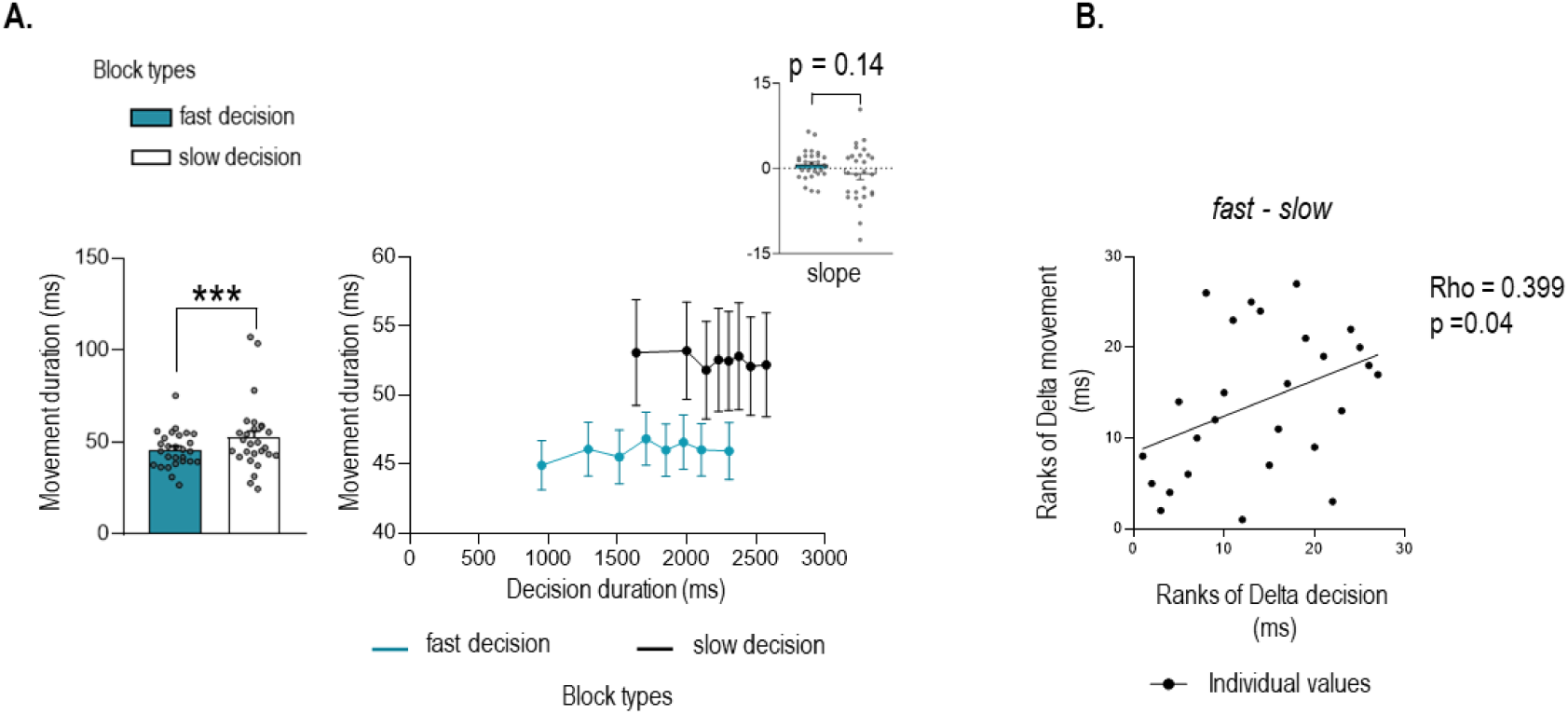
Control analyses on behavior in Experiment 1 (n=27). A. Movement duration. The results showed faster movements in fast decision (blue bar) relative to slow decision (white bar) blocks (left panel). The duration of movements as a function of decision duration evolved differently between the two types of blocks (right panel). Although the slopes did not differ from 0 for both fast and slow decision blocks (see inset), the slope for slow decision blocks still tended to be lower. **B. Correlations.** Given that deltas were computed by measuring the difference in duration between fast and slow conditions, the more the subjects showed an effect of the condition (or coregulation), the more they displayed a negative delta, which corresponds to a low rank value. So, here you can see that the more participants responded faster in fast decision blocks compare to slow ones (i.e., the smaller the rank value), the more they accelerated their movement (i.e., the smaller the rank value) in this block type. Error bars represent SE. #: t-test against 0 p < 0.025. ***p < 0.001.

**Figure S7.**
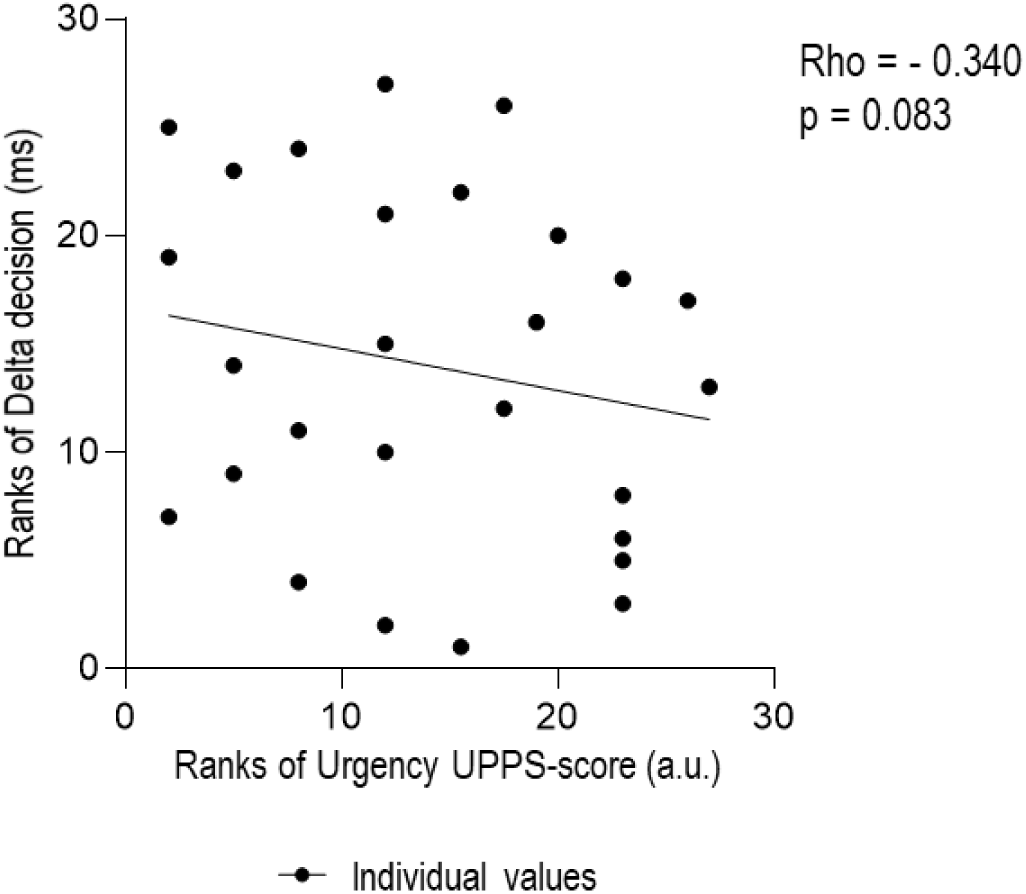
Correlation with UPPS-score in slow decision group of Experiment 2. Given that deltas were computed by measuring the difference in duration between fast and slow conditions, the more the subjects showed an effect of the condition (or tradeoff), the more they displayed a positive delta, which corresponds to a high rank value. Subjects who were more impulsive (higher rank value) tended to show less ability to slow down (lower rank value).

**Table S1.** Demographic description of fast decision and slow decision groups. The table includes the Bayesian independent t-tests for the sex, the age and the five facets of trait impulsivity of the short-UPPS. BF_10_ grades the strength of evidence for the alternative hypothesis against the null hypothesis. The error percentage (Error %) indicates the numeric robustness of the results, with low values of the Error % corresponding to a greater numerical stability of results. The gender of the participants was transformed into a numerical score to compare the average between the two groups (i.e., 1 for female (F) and 0 for male (M)).

|  | *Fast decision* | *Slow decision* | *BF<sub>10</sub>* | *Error %* |
| --- | --- | --- | --- | --- |
| *gender (M/F)* | 12/15 | 10/17 | 0.309 | 0.008 |
| *Age* | 25.11 ± 3.65 | 23.81 ± 2.87 | 0.651 | 0.008 |
| *Urgency* | 8.52 ± 3.19 | 8.37 ± 3.20 | 0.277 | 0.008 |
| *Positive urgency* | 10.63 ± 2.82 | 10.56 ± 1.85 | 0.275 | 0.008 |
| *Lack of Perseverance* | 7.85 ± 2.52 | 7.30 ± 2.45 | 0.361 | 0.008 |
| *Lack of Premeditation* | 6.70 ± 2.46 | 7.33 ± 2.45 | 0.396 | 0.008 |
| *Sensation seeking* | 11.07 ± 2.05 | 10.33 ± 3.06 | 0.430 | 0.008 |

**Table S2.** Paired Student’s t-tests of urgency level over decision duration. The critical t-value, the p-value and the Cohen’s d as a measure of the effect size are represented for each bin of decision duration (Bin_DD_). Significant p-value (with a Bonferroni-corrected threshold of .05/8) are highlighted in bold and blue.

| *Bin<sub>DD</sub>* | *t-value* | *df* | *p-value* | *Cohen's d* |
| --- | --- | --- | --- | --- |
| 1 | 12.784 | 55 | < 0.001 | 1.708 |
| 2 | 10.122 | 55 | < 0.001 | 1.353 |
| 3 | 8.298 | 55 | < 0.001 | 1.109 |
| 4 | 7.261 | 55 | < 0.001 | 0.970 |
| 5 | 6.421 | 55 | < 0.001 | 0.858 |
| 6 | 5.676 | 55 | < 0.001 | 0.758 |
| 7 | 4.816 | 55 | < 0.001 | 0.644 |
| 8 | 3.761 | 55 | < 0.001 | 0.503 |

**Table S3.** Paired Student’s t-tests of pupil dilation over decision duration. The critical t-value, the p-value and the Cohen’s d as a measure of the effect size are represented for each bin of decision duration (Bin_DD_). Significant p-value (with a Bonferroni-corrected threshold of .05/8) are highlighted in bold and blue.

| *Bin<sub>DD</sub>* | *t-value* | *df* | *p-value* | *Cohen's d* |
| --- | --- | --- | --- | --- |
| 1 | -3.086 | 17 | 0.007 | -0.727 |
| 2 | -4.007 | 17 | < 0.001 | -0.944 |
| 3 | -2.352 | 17 | 0.031 | -0.554 |
| 4 | -3.561 | 17 | 0.002 | -0.839 |
| 5 | -4.408 | 17 | < 0.001 | -1.039 |
| 6 | -3.884 | 17 | 0.001 | -0.915 |
| 7 | -3.191 | 17 | 0.005 | -0.752 |
| 8 | -5.583 | 17 | < 0.001 | -1.316 |

## References

Aron, A. R., Herz, D. M., Brown, P., Forstmann, B. U., & Zaghloul, K. (2016). Frontosubthalamic Circuits for Control of Action and Cognition. The Journal of Neuroscience, 36(45), 11489-11495. 10.1523/JNEUROSCI.2348-16.2016

Berridge, K. C., & Robinson, T. E. (2016). Liking, wanting, and the incentive-sensitization theory of addiction. American Psychologist, 71(8), 670-679. 10.1037/amp0000059

Bourgeois, A., Chelazzi, L., & Vuilleumier, P. (2016). Chapter 14—How motivation and reward learning modulate selective attention. In B. Studer & S. Knecht (Éds.), Progress in Brain Research (Vol. 229, p. 325-342). Elsevier. 10.1016/bs.pbr.2016.06.004

Carland, M. A., Thura, D., & Cisek, P. (2019). The Urge to Decide and Act : Implications for Brain Function and Dysfunction. The Neuroscientist, 25(5), 491-511. 10.1177/1073858419841553

Carsten, T., Fievez, F., & Duque, J. (2023). Movement characteristics impact decision-making and vice versa. Scientific Reports, 13(1), 3281. 10.1038/s41598-023-30325-4

Cavanagh, J. F., Wiecki, T. V., Cohen, M. X., Figueroa, C. M., Samanta, J., Sherman, S. J., & Frank, M. J. (2011). Subthalamic nucleus stimulation reverses mediofrontal influence over decision threshold. Nature Neuroscience, 14(11), 1462-1467. 10.1038/nn.2925

Charnov, E. L. (1976). Optimal foraging, the marginal value theorem. Theoretical Population Biology, 9(2), 129-136. 10.1016/0040-5809(76)90040-X

Chen, X., & Yang, T. (2021). A neural network model of basal ganglia’s decision-making circuitry. Cognitive Neurodynamics, 15(1), 17-26. 10.1007/s11571-020-09609-2

Churchland, A. K., Kiani, R., & Shadlen, M. N. (2008). Decision-making with multiple alternatives. Nature Neuroscience, 11(6), 693-702. 10.1038/nn.2123

Cisek, P., Puskas, G. A., & El-Murr, S. (2009). Decisions in Changing Conditions : The Urgency-Gating Model. The Journal of Neuroscience, 29(37), 11560-11571. 10.1523/JNEUROSCI.1844-09.2009

Cisek, P., & Thura, D. (2022). Models of Decision-Making Over Time. In P. Cisek & D. Thura, Oxford Research Encyclopedia of Neuroscience. Oxford University Press. 10.1093/acrefore/9780190264086.013.346

Coddington, L. T., & Dudman, J. T. (2019). Learning from Action : Reconsidering Movement Signaling in Midbrain Dopamine Neuron Activity. Neuron, 104(1), 63-77. 10.1016/j.neuron.2019.08.036

Derosiere, G., Thura, D., Cisek, P., & Duque, J. (2019). Motor cortex disruption delays motor processes but not deliberation about action choices. Journal of Neurophysiology, 122(4), 1566-1577. 10.1152/jn.00163.2019

Derosiere, G., Thura, D., Cisek, P., & Duque, J. (2021). Trading accuracy for speed over the course of a decision. Journal of Neurophysiology, 126(2), 361-372. 10.1152/jn.00038.2021

Derosiere, G., Thura, D., Cisek, P., & Duque, J. (2022). Hasty sensorimotor decisions rely on an overlap of broad and selective changes in motor activity. PLOS Biology, 20(4), e3001598. 10.1371/journal.pbio.3001598

Desmurget, M., & Turner, R. S. (2010). Motor Sequences and the Basal Ganglia : Kinematics, Not Habits. The Journal of Neuroscience, 30(22), 7685. 10.1523/JNEUROSCI.0163-10.2010

Ditterich, J. (2006). Evidence for time-variant decision making. European Journal of Neuroscience, 24(12), 3628-3641. 10.1111/j.1460-9568.2006.05221.x

Drugowitsch, J., Moreno-Bote, R., Churchland, A. K., Shadlen, M. N., & Pouget, A. (2012). The Cost of Accumulating Evidence in Perceptual Decision Making. The Journal of Neuroscience, 32(11), 3612-3628. 10.1523/JNEUROSCI.4010-11.2012

Dudman, J. T., & Krakauer, J. W. (2016). The basal ganglia : From motor commands to the control of vigor. Current Opinion in Neurobiology, 37, 158-166. 10.1016/j.conb.2016.02.005

Duque, J., Greenhouse, I., Labruna, L., & Ivry, R. B. (2017). Physiological Markers of Motor Inhibition during Human Behavior. Trends in Neurosciences, 40(4), 219-236. 10.1016/j.tins.2017.02.006

Duque, J., Petitjean, C., & Swinnen, S. P. (2016). Effect of Aging on Motor Inhibition during Action Preparation under Sensory Conflict. Frontiers in Aging Neuroscience, 8. https://www.frontiersin.org/articles/10.3389/fnagi.2016.00322

Eben, C., Billieux, J., & Verbruggen, F. (2020). Clarifying the Role of Negative Emotions in the Origin and Control of Impulsive Actions. Psychologica Belgica, 60(1), 1-17. 10.5334/pb.502

Ferrucci, L., Genovesio, A., & Marcos, E. (2021). The importance of urgency in decision making based on dynamic information. PLOS Computational Biology, 17(10), e1009455. 10.1371/journal.pcbi.1009455

Fievez, F., Derosiere, G., Verbruggen, F., & Duque, J. (2022). Post-error Slowing Reflects the Joint Impact of Adaptive and Maladaptive Processes During Decision Making. Frontiers in Human Neuroscience, 16, 864590. 10.3389/fnhum.2022.864590

Forstmann, B. U., van den Wildenberg, W. P. M., & Ridderinkhof, K. R. (2008). Neural Mechanisms, Temporal Dynamics, and Individual Differences in Interference Control. Journal of Cognitive Neuroscience, 20(10), 1854-1865. 10.1162/jocn.2008.20122

Frank, M. J., Samanta, J., Moustafa, A. A., & Sherman, S. J. (2007). Hold Your Horses : Impulsivity, Deep Brain Stimulation, and Medication in Parkinsonism. Science, 318(5854), 1309-1312. 10.1126/science.1146157

Gold, J. I., & Shadlen, M. N. (2007). The Neural Basis of Decision Making. Annual Review of Neuroscience, 30(1), 535-574. 10.1146/annurev.neuro.29.051605.113038

Greenhouse, I. (2022). Inhibition for gain modulation in the motor system. Experimental Brain Research, 240(5), 1295-1302. 10.1007/s00221-022-06351-5

Gross, M. P., & Dobbins, I. G. (2021). Pupil dilation during memory encoding reflects time pressure rather than depth of processing. *Journal of Experimental Psychology: Learning*, Memory, and Cognition, 47(2), 264-281. 10.1037/xlm0000818

Hanks, T., Kiani, R., & Shadlen, M. N. (2014). A neural mechanism of speed-accuracy tradeoff in macaque area LIP. eLife, 3, e02260. 10.7554/eLife.02260

Herz, D. M., Tan, H., Brittain, J.-S., Fischer, P., Cheeran, B., Green, A. L., FitzGerald, J., Aziz, T. Z., Ashkan, K., Little, S., Foltynie, T., Limousin, P., Zrinzo, L., Bogacz, R., & Brown, P. (2017). Distinct mechanisms mediate speed-accuracy adjustments in cortico-subthalamic networks. eLife, 6, e21481. 10.7554/eLife.21481

Hilz, M. J. (2022). Transcutaneous vagus nerve stimulation—A brief introduction and overview. Autonomic Neuroscience: Basic and Clinical, 243. 10.1016/j.autneu.2022.103038

Kaduk, K., Henry, T., Guitton, J., Meunier, M., Thura, D., & Hadj-Bouziane, F. (2023). Atomoxetine and reward size equally improve task engagement and perceptual decisions but differently affect movement execution. Neuropharmacology, 241, 109736. 10.1016/j.neuropharm.2023.109736

Kelly, S. P., Corbett, E. A., & O’Connell, R. G. (2021). Neurocomputational mechanisms of prior-informed perceptual decision-making in humans. Nature Human Behaviour, 5(4), 467-481.

Kita, K., Du, Y., & Haith, A. M. (2023). Evidence for a common mechanism supporting invigoration of action selection and action execution. Journal of neurophysiology, 130(2), 238–246. 10.1152/jn.00510.2022

Klein, P.-A., Petitjean, C., Olivier, E., & Duque, J. (2014). Top-down suppression of incompatible motor activations during response selection under conflict. NeuroImage, 86, 138-149. 10.1016/j.neuroimage.2013.08.005

Lawlor, J., Zagala, A., Jamali, S., & Boubenec, Y. (2023). Pupillary dynamics reflect the impact of temporal expectation on detection strategy. iScience, 26(2), 106000. 10.1016/j.isci.2023.106000

Lemon, W. C. (1991). Fitness consequences of foraging behaviour in the zebra finch. Nature, 352(6331), 153-155. 10.1038/352153a0

Mazzoni, P., Hristova, A., & Krakauer, J. W. (2007). Why Don’t We Move FasterParkinson’s Disease, Movement Vigor, and Implicit Motivation. The Journal of Neuroscience, 27(27), 7105. 10.1523/JNEUROSCI.0264-07.2007

McGinley, M. J., David, S. V., & McCormick, D. A. (2015). Cortical Membrane Potential Signature of Optimal States for Sensory Signal Detection. Neuron, 87(1), 179-192. 10.1016/j.neuron.2015.05.038

Mosher, C. P., Mamelak, A. N., Malekmohammadi, M., Pouratian, N., & Rutishauser, U. (2021). Distinct roles of dorsal and ventral subthalamic neurons in action selection and cancellation. Neuron, 109(5), 869–881.e6. 10.1016/j.neuron.2020.12.025

Murphy, P. R., Boonstra, E., & Nieuwenhuis, S. (2016). Global gain modulation generates time-dependent urgency during perceptual choice in humans. Nature Communications, 7(1), 13526. 10.1038/ncomms13526

Murphy, P. R., Robertson, I. H., Balsters, J. H., & O’connell, R. G. (2011). Pupillometry and P3 index the locus coeruleus–noradrenergic arousal function in humans. Psychophysiology, 48(11), 1532-1543. 10.1111/j.1469-8986.2011.01226.x

Niv, Y. (2007). Cost, Benefit, Tonic, Phasic : What Do Response Rates Tell Us about Dopamine and Motivation? Annals of the New York Academy of Sciences, 1104(1), 357-376. 10.1196/annals.1390.018

O’Connell, R. G., Shadlen, M. N., Wong-Lin, K., & Kelly, S. P. (2018). Bridging Neural and Computational Viewpoints on Perceptual Decision-Making. Trends in Neurosciences, 41(11), 838-852. 10.1016/j.tins.2018.06.005

Oldfield, R. C. (1971). The assessment and analysis of handedness : The Edinburgh inventory. Neuropsychologia, 9(1), 97-113. 10.1016/0028-3932(71)90067-4

Park, J., Coddington, L. T., & Dudman, J. T. (2020). Basal Ganglia Circuits for Action Specification. Annual Review of Neuroscience, 43(1), 485-507. 10.1146/annurev-neuro-070918-050452

Quoilin, C., Fievez, F., & Duque, J. (2019). Preparatory inhibition : Impact of choice in reaction time tasks. Neuropsychologia, 129, 212-222. 10.1016/j.neuropsychologia.2019.04.016

Ratcliff, R., & Smith, P. L. (2004). A Comparison of Sequential Sampling Models for Two-Choice Reaction Time. Psychological Review, 111(2), 333-367. 10.1037/0033-295X.111.2.333

Ratcliff, R., Smith, P. L., Brown, S. D., & McKoon, G. (2016). Diffusion Decision Model : Current Issues and History. Trends in Cognitive Sciences, 20(4), 260-281. 10.1016/j.tics.2016.01.007

Reppert, T. R., Heitz, R. P., & Schall, J. D. (2023). Neural mechanisms for executive control of speed-accuracy trade-off. Cell Reports, 42(11), 113422. 10.1016/j.celrep.2023.113422

Reynaud, A. J., Saleri Lunazzi, C., & Thura, D. (2020). Humans sacrifice decision-making for action execution when a demanding control of movement is required. Journal of Neurophysiology, 124(2), 497-509. 10.1152/jn.00220.2020

Şahіn, M., & Aybek, E. (2020). Jamovi : An Easy to Use Statistical Software for the Social Scientists. International Journal of Assessment Tools in Education, 6(4), 670-692. 10.21449/ijate.661803

Salamone, J. D., & Correa, M. (2024). The Neurobiology of Activational Aspects of Motivation : Exertion of Effort, Effort-Based Decision Making, and the Role of Dopamine. Annual Review of Psychology, 75(1), annurev-psych-020223-012208. 10.1146/annurev-psych-020223-012208

Saleri Lunazzi, C., Reynaud, A. J., & Thura, D. (2021). Dissociating the Impact of Movement Time and Energy Costs on Decision-Making and Action Initiation in Humans. Frontiers in Human Neuroscience, 15. 10.3389/fnhum.2021.715212

Schultz, W., Stauffer, W. R., & Lak, A. (2017). The phasic dopamine signal maturing : From reward via behavioural activation to formal economic utility. Current Opinion in Neurobiology, 43, 139-148. 10.1016/j.conb.2017.03.013

Shadmehr, R., Huang, H. J., & Ahmed, A. A. (2016). A Representation of Effort in Decision-Making and Motor Control. Current Biology, 26(14), 1929-1934. 10.1016/j.cub.2016.05.065

Shadmehr, R., Reppert, T. R., Summerside, E. M., Yoon, T., & Ahmed, A. A. (2019). Movement Vigor as a Reflection of Subjective Economic Utility. Trends in Neurosciences, 42(5), 323-336. 10.1016/j.tins.2019.02.003

Spieser, L., Servant, M., Hasbroucq, T., & Burle, B. (2017). Beyond decision ! Motor contribution to speed–accuracy trade-off in decision-making. Psychonomic Bulletin & Review, 24(3), 950-956. 10.3758/s13423-016-1172-9

Steinemann, N. A., O’Connell, R. G., & Kelly, S. P. (2018). Decisions are expedited through multiple neural adjustments spanning the sensorimotor hierarchy. Nature Communications, 9(1), 3627. 10.1038/s41467-018-06117-0

Steudel-Numbers, K. L., & Wall-Scheffler, C. M. (2009). Optimal running speed and the evolution of hominin hunting strategies. Journal of Human Evolution, 56(4), 355-360. 10.1016/j.jhevol.2008.11.002

Thura, D. (2020). Decision urgency invigorates movement in humans. Behavioural Brain Research, 382, 112477. 10.1016/j.bbr.2020.112477

Thura, D., Beauregard-Racine, J., Fradet, C.-W., & Cisek, P. (2012). Decision making by urgency gating : Theory and experimental support. Journal of Neurophysiology, 108(11), 2912-2930. 10.1152/jn.01071.2011

Thura, D., Cabana, J.-F., Feghaly, A., & Cisek, P. (2022). Integrated neural dynamics of sensorimotor decisions and actions. PLOS Biology, 20(12), e3001861. 10.1371/journal.pbio.3001861

Thura, D., & Cisek, P. (2017). The Basal Ganglia Do Not Select Reach Targets but Control the Urgency of Commitment. Neuron, 95(5), 1160–1170.e5. 10.1016/j.neuron.2017.07.039

Thura, D., Cos, I., Trung, J., & Cisek, P. (2014). Context-Dependent Urgency Influences Speed–Accuracy Trade-Offs in Decision-Making and Movement Execution. The Journal of Neuroscience, 34(49), 16442. 10.1523/JNEUROSCI.0162-14.2014

Vassiliadis, P., Derosiere, G., Grandjean, J., & Duque, J. (2020). Motor training strengthens corticospinal suppression during movement preparation. Journal of Neurophysiology, 124(6), 1656-1666. 10.1152/jn.00378.2020

Wagenmakers, E.-J., Marsman, M., Jamil, T., Ly, A., Verhagen, J., Love, J., Selker, R., Gronau, Q. F., Šmíra, M., Epskamp, S., Matzke, D., Rouder, J. N., & Morey, R. D. (2018). Bayesian inference for psychology. Part I: Theoretical advantages and practical ramifications. Psychonomic Bulletin & Review, 25(1), 35-57. 10.3758/s13423-017-1343-3

Weinstein, A. M. (2023). Reward, motivation and brain imaging in human healthy participants – A narrative review. Frontiers in Behavioral Neuroscience, 17, 1123733. 10.3389/fnbeh.2023.1123733

Wessel, J. R., & Aron, A. R. (2017). On the Globality of Motor Suppression : Unexpected Events and Their Influence on Behavior and Cognition. Neuron, 93(2), 259-280. 10.1016/j.neuron.2016.12.013

Wilhelm, E., Quoilin, C., Derosiere, G., Paço, S., Jeanjean, A., & Duque, J. (2022). Corticospinal Suppression Underlying Intact Movement Preparation Fades in Parkinson’s Disease. Movement Disorders, 37(12), 2396-2406. 10.1002/mds.29214

Yoon, T., Geary, R. B., Ahmed, A. A., & Shadmehr, R. (2018). Control of movement vigor and decision making during foraging. Proceedings of the National Academy of Sciences, 115(44). 10.1073/pnas.1812979115

Yttri, E. A., & Dudman, J. T. (2018). A Proposed Circuit Computation in Basal Ganglia : History-Dependent Gain. Movement Disorders, 33(5), 704-716. 10.1002/mds.27321

Zarrugh, M. Y., Todd, F. N., & Ralston, H. J. (1974). Optimization of energy expenditure during level walking. European Journal of Applied Physiology and Occupational Physiology, 33(4), 293-306. 10.1007/BF00430237

Zénon, A. (2019). Eye pupil signals information gain. Proceedings of the Royal Society B: Biological Sciences, 286(1911), 20191593. 10.1098/rspb.2019.1593

